# Split-miniSOG for detecting and localizing intracellular protein-protein interactions: application to correlated light and electron microscopy

**DOI:** 10.1101/423566

**Authors:** Daniela Boassa, Sakina P. Lemieux, Varda Lev-Ram, Junru Hu, Qing Xiong, Sebastien Phan, Mason Mackey, Ranjan Ramachandra, Stephen R. Adams, Roger Y. Tsien, Mark H. Ellisman, John T. Ngo

## Abstract

A protein complementation assay (PCA) for detecting and localizing intracellular protein-protein interactions (PPIs) was built by bisection of miniSOG, a fluorescent flavoprotein derived from the light, oxygen, voltage (LOV)-2 domain of *Arabidopsis* phototropin. When brought together by interacting proteins, the fragments reconstitute a functional reporter that permits tagged protein complexes to be visualized by fluorescence light microscopy (LM), and then by standard as well as “multicolor” electron microscopy (EM) imaging methods via the photooxidation of 3-3’-diaminobenzidine (DAB) and its lanthanide-conjugated derivatives.

---

PPIs underlie countless cellular processes and identifying when and where these interactions occur is essential for understanding their roles in biology and disease. Given their critical roles, a variety of different tools have been developed in order to track and quantify PPIs, including tools monitoring their formation and dynamics, as well as probes for visualizing their spatial distribution within intact cells^1^. However, what remains missing are facile methods for visualizing PPIs in high resolution, which is important to do given that PPIs often give rise to protein complexes that localize to specific cellular microdomains. Methods for imaging PPIs with sub-diffraction resolution would represent powerful tools, as biomacromolecules often interact to rise to nanometer-scale assemblies.

Here, we describe the development of a novel PCA that can be used to track PPIs by correlative light and electron microscopy (CLEM). In this approach, PPI-partners are tagged with split-protein fragments derived from miniSOG, a fluorescent flavoprotein and genetically encoded EM-compatible tag engineered from the LOV2 domain of *Arabidopsis* phototropin^2^. Due to the mutation of LOV2’s adduct-forming cysteine to glycine, miniSOG is both green-fluorescent and able to generate reactive oxygen species upon illumination with blue light. In fixed cells, photogenerated singlet oxygen (^1^O_2_) from miniSOG-tagged proteins can be used to convert 3,3’-diaminobenzidine (DAB) into a highly localized polymeric precipitate, which upon subsequent reaction with osmium tetroxide (OsO_4_), is readily identified under the electron microscope.

Reasoning that a split-version of miniSOG could be applied to image PPIs by CLEM in a manner similar to how split-GFP is used in fluorescence microscopy, we set out to identify a site in the LOV domain sequence where it could be bisected into PCA-suitable fragments. Given that the solubility of proteins has previously been found to influence their “split-ability” (such as in the case of the “super-folder GFP,” which can be split at multiple positions), we generated a more soluble version of miniSOG by fusing it to LOV2’s native Jα-helix (miniSOG-Jα, **Fig 1**). NMR and x-ray crystallographic studies have shown that the amphiphilic Jα-helix docks to LOV2 through a hydrophobic face while orienting a hydrophilic face toward the protein surface^3-5^. Given these favorable interactions, we anticipated that the fusion of Jα-helix would enhance the stability of miniSOG, and in turn favor the identification of a functional split-site. Indeed, miniSOG-Jα exhibited a reduced propensity to form inclusion bodies when overexpressed in *E. coli* as compared to the original miniSOG (**Supplementary Fig. 1**).

**Figure 1.**
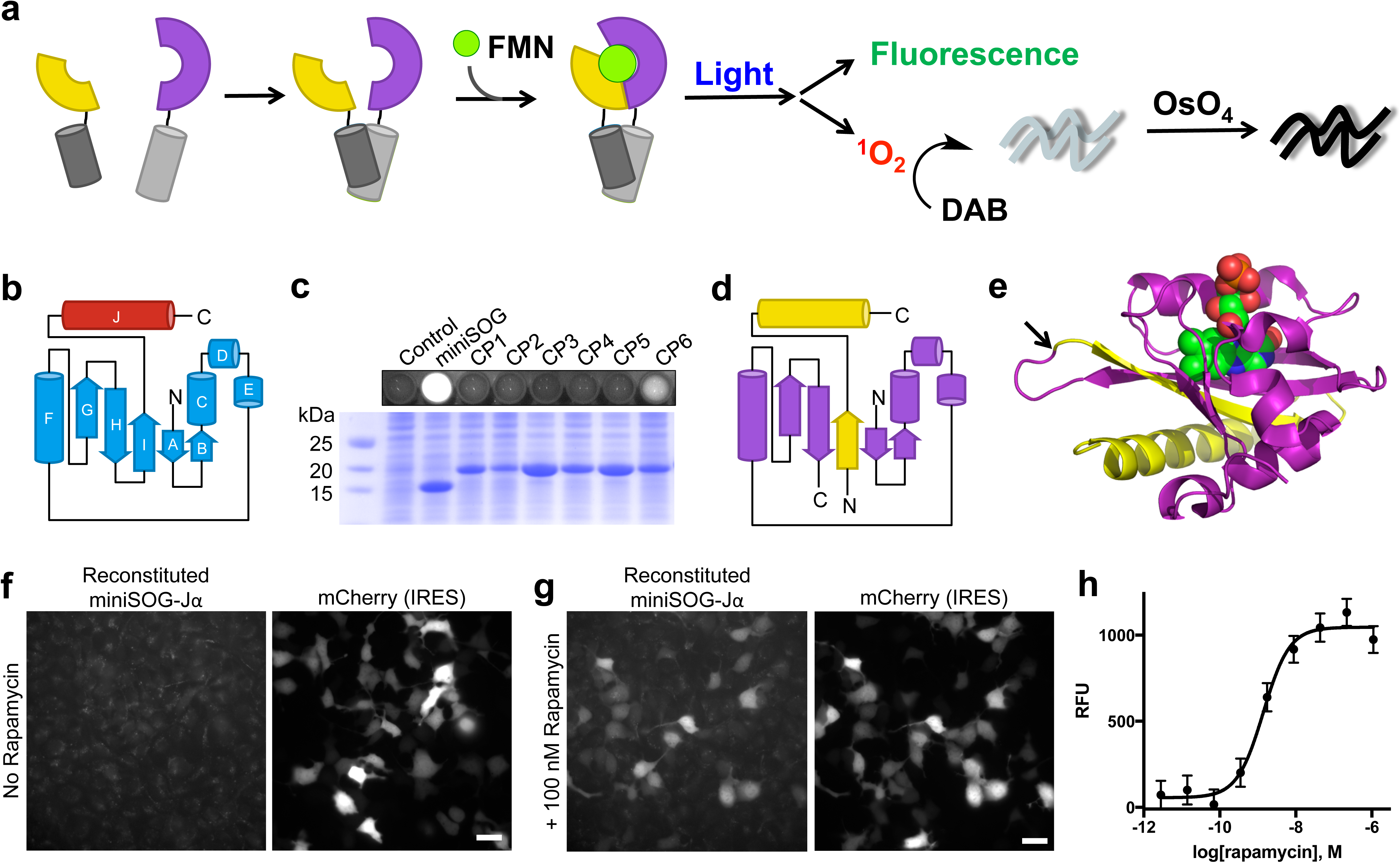
Identification of a functional bisection site for generating PCA-suitable miniSOG-Jα fragments. (**a**) Schematic depicting miniSOG complementation. MiniSOG is divided into fragments (yellow and purple) that do not bind FMN (green) unless brought together by interacting proteins (dark and light gray cylinders). Reconstituted complexes can be imaged by fluorescence and correlated to EM observations via photogeneration of ^1^O_2_ for localized polymerization of DAB. (**b**) Topology of miniSOG-Jα with miniSOG shown in blue and the Jα-helix in red. (**c**) Fluorescence from *E. coli* cells expressing various CPs (top) and a Coomassie-stained SDS-PAGE gel of corresponding cell lysates (bottom). (**d**) Topology of mSOG_1-94_ (purple) and mSOG-Jα_95-140_ (yellow), polypeptide fragments resulting from bisection of miniSOG-Jα between Gly94 and Glu95. (**e**) Structure of the related *A. sativa* LOV2 domain (PDB: 2V0U) showing FMN (spheres), as well as residues corresponding to mSOG_1-94_ (purple) and mSOG-Jα_95-140_ (yellow). (**f-g**) Fluorescence imaging of HEK293 cells co-transfected with mSOG_1-94_-FRB and FKBP-mSOG-Jα_95-140_. Signal from reconstituted miniSOG-Jα was detected only in rapamycin-treated cells. Transfected cells were identified by IRES-mediated expression of mCherry. Scale bars, 20 microns. (**h**) Quantification of rapamycin-induced miniSOG-Jα fluorescence. The plotted values represent geometric means, mean ± s.d. of n=3 biologically independent samples.

Given that viable bisection and circular permutation points often coincide^6^, we next sought to identify viable circular permutants (CPs) of miniSOG-Jα circular, which we anticipated would guide toward a viable protein split-site. In an initial pool, we generated candidate CPs by introducing new termini within miniSOG-Jα loop regions containing three or more amino acids (**Supplementary Fig. 2**). Rationalizing that a properly folded CP would retain the ability to bind FMN, the proteins were expressed in bacteria and evaluated on the basis of cellular fluorescence (**Fig. 1c**). From a set of six CPs, we identified a functional sequence termed “CP6,” corresponding to circular permutation of miniSOG-Jα at Glu95. When compared to a sequence lacking the Jα segment (CP6ΔJα), CP6 let to substantially higher cellular fluorescence (**Supplementary Fig. 3**), and the measurements using purified CP6 indicated a fluorescence quantum yield that was comparable to that of the parental miniSOG (**Supplementary Table 1**).

We next bisected miniSOG-Jα between Gly94 and Glu95 (**Supplementary Fig. 4**), producing an N-terminal segment termed “mSOG_1-94_” (10.96kDa) and a 46 amino acid C-terminal polypeptide designated “mSOG-Jα_95-140_” (5.15 kDa), which we evaluated for use in a PCA. First, using complementary leucine zipper sequences tagged with each fragment, we tested their ability to reconstitute a FMN-binding domain in *E. coli*. Indeed, cells that co-expressed the tagged domains exhibited fluorescence emission corresponding to miniSOG-bound chromophore (**Supplementary Fig. 5, 6**).

Furthermore, His-tag purification resulted in a chromophore-bound protein complexes with spectral properties characteristic to LOV-bound FMN, and a fluorescence quantum yield similar to that of miniSOG (**Supplementary Table 1**, ΦF = 0.33).

To test whether the fragments could complement one another in mammalian cells, and to determine whether the reconstitution of miniSOG-Jα required physical interaction between tagged domains, we fused mSOG_1-94_ and mSOG-Jα_95-140_ to domains that undergo rapamycin-inducible interaction^7^. Using HEK293 cells that co-expressed proteins, we quantified the extent of miniSOG-Jα fluorescence in cells treated with increasing concentrations of rapamycin (**Fig 1f-h**). In analyses by flow cytometry and fluorescence microscopy, drug-untreated cells exhibited only background emission levels, comparing to that of non-transfected controls. However, cells treated with rapamycin exhibited fluorescence intensities corresponding to a dose-dependent reconstitution of miniSOG-Jα. Indeed, these results confirmed that complementation between mSOG_1-94_ and mSOG-Jα_95-140_ occurred in a PPI-dependent manner.

To determine whether miniSOG-Jα complementation could be used to image PPIs via CLEM, we fused mSOG_1-94_ and mSOG-Jα_95-140_ to the basic region-leucine zipper domains of bFos and bJun, subunits of the AP-1 transcriptional complex that interact constitutively as nuclear heterodimers^8^. Co-expression of the tagged zippers in mammalian cells resulted in bright nuclear fluorescence with enrichment in nucleoli (**Fig. 2a**), consistent with the known localization of bFos/bJun complexes as previously determined using split-YFP^9^. To test whether the tagged complexes could be visualized by EM, co-transfected HeLa cells were photooxidized using DAB (see Online Methods). Following photooxidation, optically-dense reaction products were visible under transmitted light in the nuclei of positive cells in patterns matching the distribution of reconstituted miniSOG-Jα fluorescence (**Fig. 2b**). At the EM level, these exhibited significantly increased nuclear contrast on electron micrographs, which was especially apparent upon comparison of expressing and non-expressing cells within the photooxidized area (**Fig. 2c-e**).

**Figure 2.**
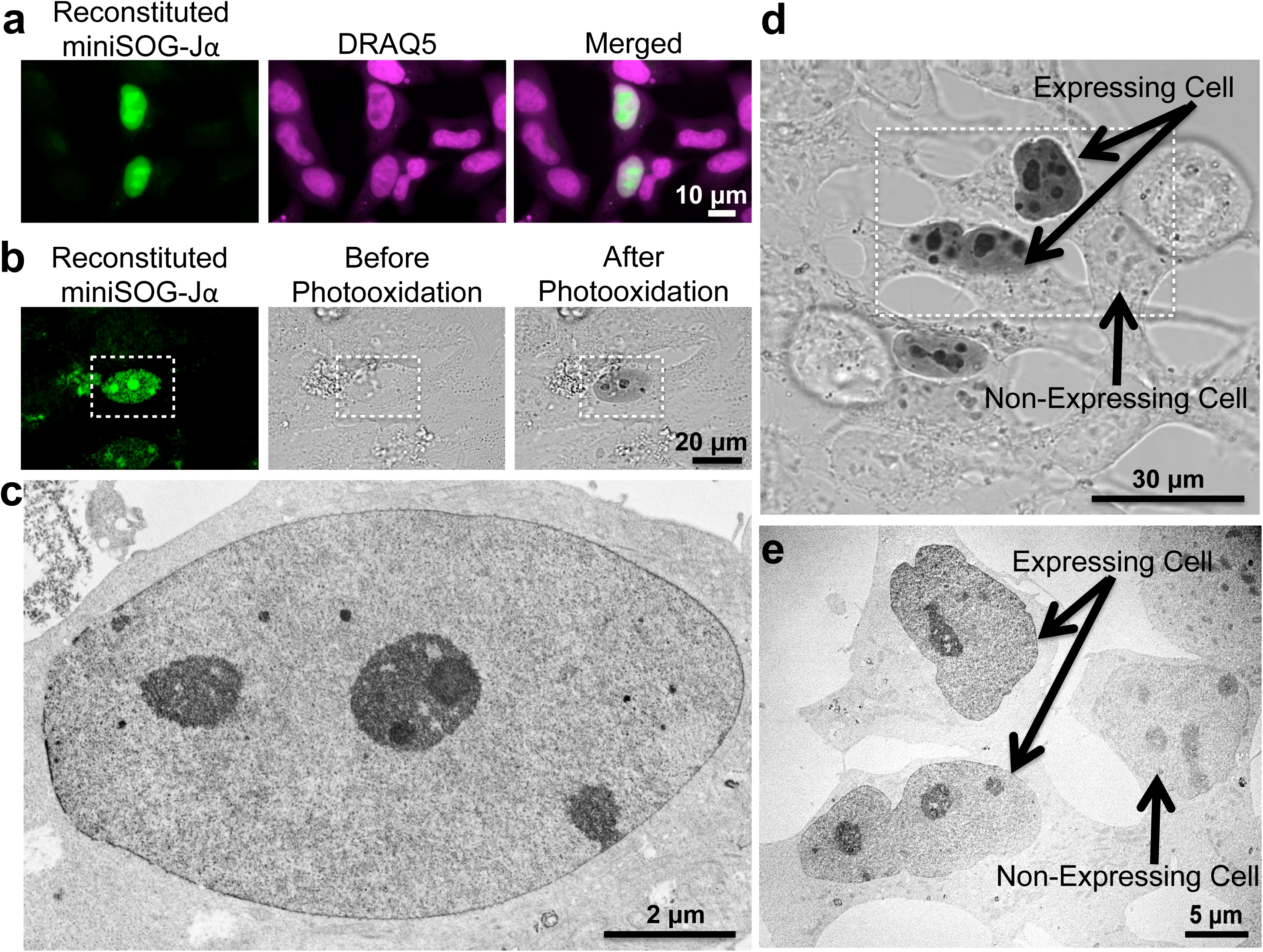
Correlated light and EM imaging of nuclear mSOG_1-94_ -bFos/bJun-mSOG-J _*α* 95-140_ complexes. **(a)** Epifluorescence images showing colocalization of reconstituted miniSOG-J *α* complexes with DRAQ5 (a far-red emitting nuclear stain) in HEK293 cells. (**b**) Co-transfected cells exhibited similar nuclear fluorescence, as revealed by confocal fluorescence imaging (left). Transmitted light images of the same area showing cells before (middle) and after (right) a 3-minute illumination with intense blue light in the presence of DAB. Following photooxidation, an optically-dense DAB reaction product was observed in the positive cell (white rectangle). (**c**) TEM micrograph of the same cell indicated in (b) showing the OsO4-stained DAB reaction products as darkened contrast. (**d**) Direct comparison of cells positive for mSOG_1-94_-bFos and bJun-mSOG-J _*α* 95-140_ (“expressing cell”) to nearby non-expressing cells by transmitted light imaging and EM (**e**, micrograph of the same area indicated by the white rectangle in **d**). The cells resided within a photooxidation area that was illuminated for 8 minutes.

Recognizing that reversibility in split-reporter systems is a desirable feature of PCAs, we tested whether the reconstituted miniSOG-Jα complex could be dissociated in the absence of a PPI template. Using a purified, single-chain construct in which a tobacco etch virus (TEV)-protease cleavage site was inserted into miniSOG-Jα between Gly94 and Glu95, we determined the state of the FMN chromophore before and after digestion with TEV protease (**Supplementary Fig. 7**). Optical and fluorescence analyses indicated that the FMN was bound to the protein prior to cleavage but was released upon digestion with TEV protease—suggesting that in the absence of a linkage, mSOG_1-94_, mSOG-Jα_95-140_ and FMN readily dissociate from one another. In addition, immunoprecipitation (IP) analyses were performed using a self-assembling protein (FM_1_, a mutant of FKBP14)^10^ that can be monomerized using a specific ligand, FK506. In the absence of FK506, IP of FM_1_-mSOG-Jα_95-140_ via a fused hemagglutinin (HA) tag resulted in the co-precipitation of FM1-mSOG_1-94_. However, treatment with FK506 to dissociate FM_1_ dimers abolished co-IP of FM1-mSOG_1-94_ (**Supplementary Fig. 8**). Together, these results indicate that the components of the reconstituted miniSOG-Jα ternary complex interact in a reversible and dynamic manner.

In order to demonstrate the utility of the probe in studying disease-relevant PPIs, we next applied split-miniSOG-Jα to visualize neurotoxic assemblies of α-synuclein (α-syn), a neuronal protein involved in Parkinson’s disease^11-12^. Using tagged monomers of α-syn, we carried out correlative analyses in order to observe the self-aggregation of the protein within neuronal cells (**Fig. 3a**). At the level of LM, we observed a diffuse fluorescent signal throughout the expressing cells with brighter labeling of α-syn aggregates accumulated in the soma and dendrites of transfected neurons (**Fig. 3b**), formations consistent with ‘Lewy bodies’ and ‘Lewy neurites’ that are observed in the post-mortem histological sections of individuals afflicted with Parkinson’s disease, and other synucleinopathies. Following DAB photooxidation and OsO4-staining, we visualized these assemblies using transmission EM as well as 3-dimensional EM tomography^13^ (**Fig. 3, 4, Supplementary Fig. 9**). In the somata and in neuronal processes, we observed darkly stained fibrillar structures and spherical assemblies (**Fig. 4**) likely representing various protofibrillar intermediates. We observed both straight and twisted α-syn filaments with a diameter ranging from 4 to 15 nm, consistent with what has been observed *in vitro* for purified protein^14^. In dendrites of expressing neurons these α-syn aggregates were contained in membrane-limited organelles, fusing to the plasma membrane suggesting release to the extracellular compartment (**Supplementary Fig. 9)**. Split-miniSOG-Jα and CLEM imaging allowed for the first time to image these structures *in vivo*.

**Figure 3.**
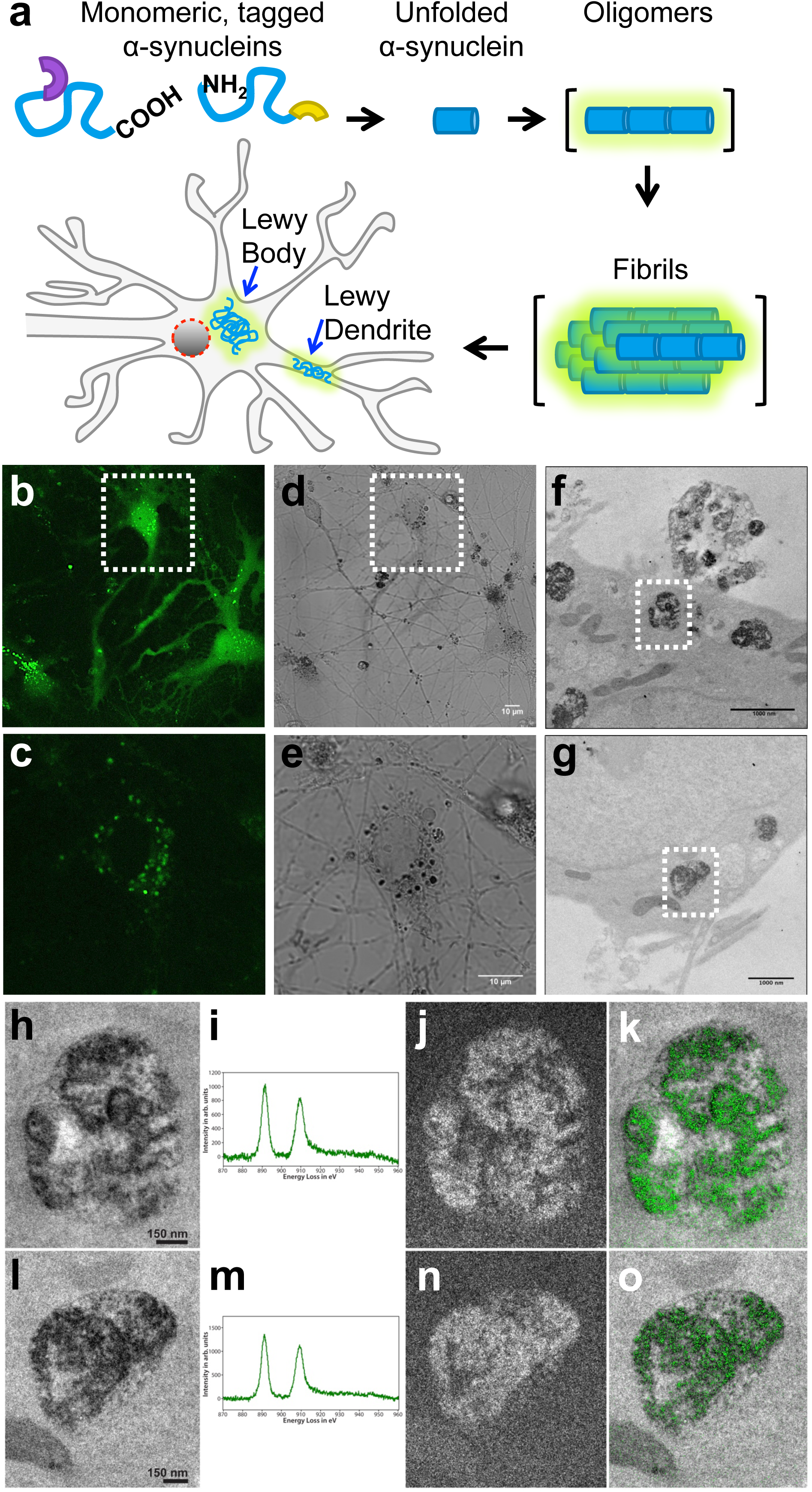
Correlated light and EM imaging of wild-type α-syn oligomers within neurons highlights an enhanced lysosomal pathology. (**a**) Schematic depiction of the chimeras used to visualize α-syn aggregates. mSOG_1-94_ (purple) and mSOG-Jα_95-140_ (yellow) were fused to the N-terminus and C-terminus of α-syn. Both N-terminus/C-terminus and C-terminus/C-terminus fusions were tested with similar results. Aggregation of the proteins leads to miniSOG-Jα reconstitution. (**b-c**) Confocal fluorescence image (single confocal plane) of live cultured neurons co-expressing the tagged proteins. A diffuse fluorescent signal is observed throughout the cells with brighter labeling of α-syn aggregates in the soma and neurites. Image in (**c**) represents the same area indicted by the white rectangle in (**b)**. (**d-e**) Corresponding transmitted light images were acquired post-fixation after 5 minutes and 30 seconds illumination in the presence of DAB. Image in (**e)** represents the same area indicted by the white rectangle in (**d)**. (**f-g**) TEM images of α-syn aggregates following DAB photooxidation, OsO_4_-staining, embedding and sectioning. Labeled α-syn aggregates were observed as darkly stained inclusions in the perinuclear region of the neuronal cell bodies. These appear to be in the lumen of lysosomes. (**h-o**) EELS and EFTEM confirm specificity of α-syn aggregates labeling. (**h**) Conventional TEM image of α-syn aggregates (corresponding to white box in (**f**)) obtained at a magnification of 15 kx (pixel size of 0.35 nm/pixel). (**i**) Electron Energy-Loss Spectra showing the presence of a strong Cerium signal within the lysosomal structures. (**j**) A Ce elemental map obtained at a magnification of 15 kx on the direct detector device DE-12. The pre-edge and post-edge images were a sum of 14 individual drift corrected images, each acquired for a 125s exposure. (**k**) A single-color merge of the elemental map (green for Ce), overlaid on the conventional image. (**l**) Conventional TEM image of α-syn aggregates (corresponding to white box in (**g**)), obtained at a magnification of 12 kx (pixel size of 0.43 nm/pixel). (**m**) Electron Energy-Loss Spectra showing the presence of a strong Cerium signal. (**n**) A Ce elemental map obtained at a magnification of 12 kx on the direct detector device DE-12. The pre-edge and post-edge images were a sum of 11 individual drift corrected images, each acquired for a 125s exposure. (**o**) A single-color merge of the elemental map (green for Ce), overlaid on the conventional image. A Gaussian smoothing blur of radius 3 pixel was applied to all the images.

**Figure 4.**
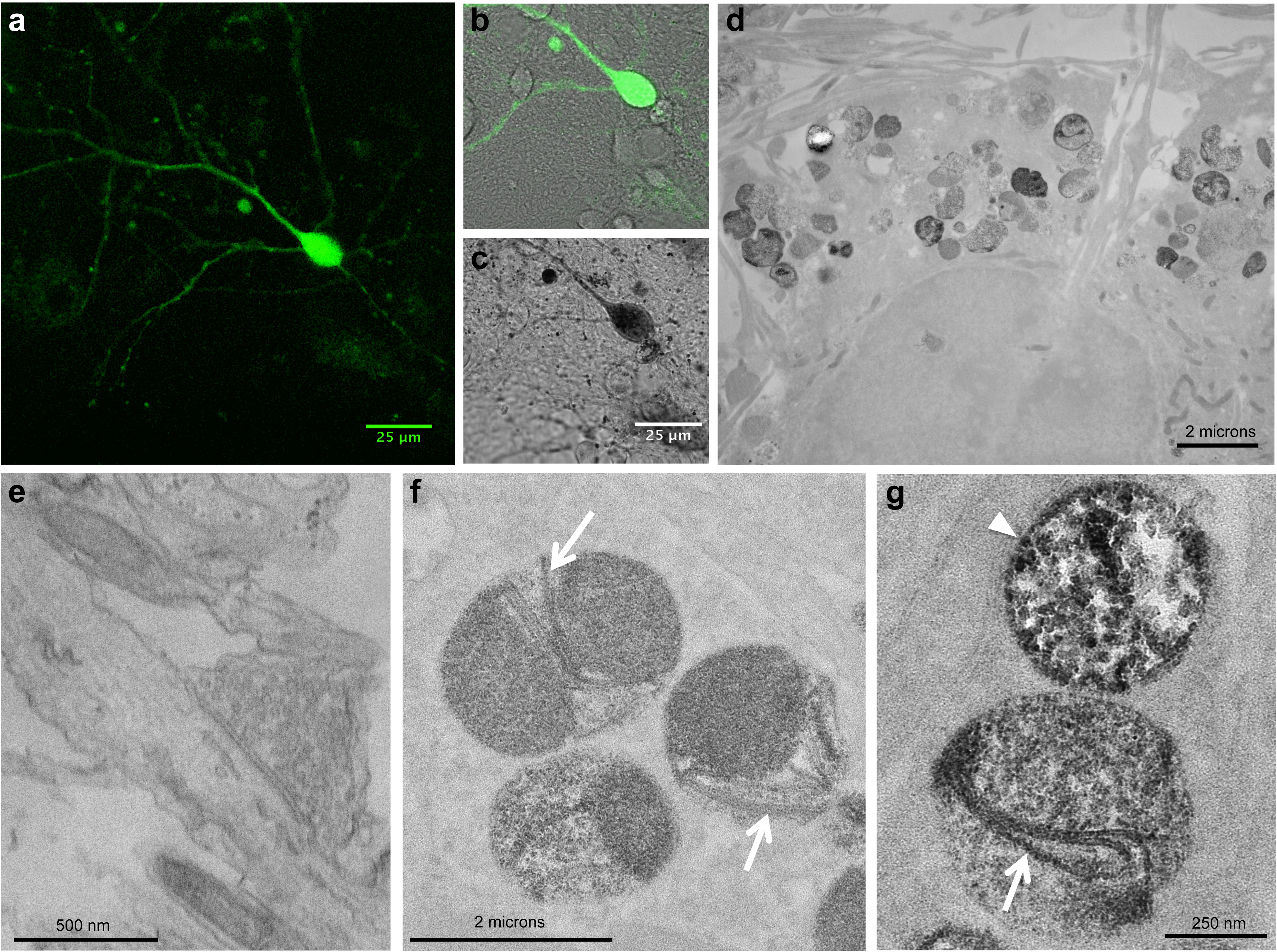
Correlated light and EM imaging of wild-type α-syn aggregates reveals fibrillar structures and protofibrillar intermediates within neurons. (**a**) Confocal fluorescence image of cultured neurons expressing α-syn aggregates. (**b**) Fluorescence image overlaid on the transmitted light image before photooxidation of DAB. (**c**) Following photooxidation, the appearance of the optically visible DAB reaction product correlates with the detected reconstituted miniSOG fluorescence in (a, b). (**d**) Electron micrograph showing labeled α-syn aggregates observed as darkly stained inclusions contained in lysosomes in the perinuclear region of the neuronal cell bodies (**d, f**), and in dendritic regions (**g**). (**e**) No labeling was observed at presynaptic terminals where α-syn aggregates were not detected. (**f, g**) At high magnification, fibrillar structures were clearly visible within these inclusions in lysosomes (white arrows), as well as spherical and chainlike protofibrillar intermediates (white arrowhead).

In addition to their presence in the soma and in neuronal processes, we also observed stained inclusions within the lumen of lysosomes. To confirm the labeling specificity of the tagged aggregates within these organelles, which are inherently electron-dense, we applied a “multicolor” EM labeling strategy (**Fig. 3h-o**) using a lanthanide-chelated analog of DAB^15^. Like DAB, these analogs are able to form localized polymeric products upon reaction with photogenerated ^1^O_2_. However, in addition to osmification, the polymeric product can also be spatially mapped using energy-filtered EM and electron energy-loss spectroscopy (EELS). Thus, the precipitated lanthanide atoms serve as an atomic signature of the polymeric labels deposited by photooxidation. This allowed us to confirm the composition of the lysosomal inclusions as α-syn aggregates. Furthermore, such labeling was not observed in control conditions (non-photooxidized expressing neurons and photooxidized untransfected neurons, **Supplementary Fig. 10**).

Fluorescence and DAB signals corresponding to reconstituted miniSOG-Jα were not detected within the axon terminals of neurons expressing tagged wild-type α-syn, nor in those containing tagged versions of the disease-associated A53T α-syn mutant (**Fig 4e**, **Supplementary Fig. 11, 12**). These observations were especially surprising given their contrast to previous data in which α-syn association was imaged via split-Venus complementation^16^. As a potential explanation for this discrepancy, we hypothesized that the Venus-positive signals observed in axon terminals were due to irreversible nature of the reconstituted split-Venus probe. In fact, while complementation of split miniSOG is reversible and does not increase avidity of the partners, the complementation of Venus is irreversible—a property that could potentially promote the artefactual aggregation of the interacting proteins by preventing their subsequent dissociation.

To confirm that the lack of axon terminal-localized signal was not due to the absence of split-miniSOG-Jα-tagged fragments at these subcellular sites, we stained transfected neurons using an antibodies against the α-syn chimeras, confirming that both the mSOG_1-94_- and mSOG-Jα_95-140_-tagged versions of the protein were localized to presynaptic sites (**Supplemental Fig. 13**). Indeed, these results indicate that the lack of miniSOG-Jα complementation appears not to be due to an inability of the tagged species to traffic to axon terminals, but rather a difference in their assembly state these subcellular positions. In addition to the split-miniSOG-Jα constructs, we also tested a single-chain fused dimer of α-syn, in which miniSOG was inserted between two copies of the self-aggregating protein (α-syn-miniSOG-α-syn). Given its single-chain nature, we anticipated that α-syn-miniSOG-α-syn would mimic the product formed upon complementation of split-Venus tagged monomers, essentially producing a permanently-associated “dimer.” Intriguingly, neurons expressing α-syn-miniSOG-α-syn exhibited fluorescence throughout the transfected cells, including intense signals that were localized to presynaptic sites and produced intense DAB reaction products upon photooxidation (**Supplemental Fig. 14**). Thus, although previously reported results using split-Venus-tagged constructs suggest that α-syn oligomers cluster at synaptic vesicles in order to restricting vesicle trafficking and recycling, our observations regarding the localization of oligomeric α-syn species suggest a different scenario into the neurotoxic role of oligomers and regarding where, for example, drugs meant to reduce oligomerization might act. Overall these results highlight the great advantages in using the split miniSOG complementation system in detecting intracellular protein-protein interactions with high spatial resolution, particularly for studies on aggregation of proteins associated with neurodegenerative diseases *in situ*.

The ability to track specified biomolecules by EM has progressed significantly since the initial introduction of miniSOG, and these developments in some ways resemble those that were made in the years following the introduction of the initial GFP. The work described herein contributes to that progress through the introduction of a simple technique for multi-scale visualization of PPIs via CLEM, enabling the imaging of intracellular complexes from the micron- to nanometer-scales. The system should complement the recently described probes for imaging extracellular and luminal PPIs^17^, as well as those that have been recently developed to visualize non-protein biomolecules by EM^18-19^. Combination of these techniques with multi-color EM strategies^15^ should provide a powerful strategy for the simultaneous detection of multiple biochemical parameters, not only in high resolution, but also within detailed ultrastructural contexts.

Future adaptations of split-miniSOG-Jα may provide routes to EM imaging of diverse biochemical events (beyond PPIs) through design of reporters that sense and align fragments in response to specific post-translational modifications or nucleic acid sequences^1^. Finally, we believe that the split site identified here will also be transferable to other LOV domain-based tools, as the bisection of the related fluorescent reporter protein iLOV^7^ at the same position was also successful (**Supplementary Fig. 15**).

## Online methods

### Sequence Information

Mammalian expression constructs encoding mSOG_1-94_ and mSOG-Jα_95-140_ can be obtained directly through Addgene.

### Comparison of miniSOG and miniSOG-Jα

DNA sequences encoding miniSOG and miniSOG-Jα were synthesized by PCR overlap-extension and inserted into a bacterial expression vector (pQE-80L, Qiagen) in-frame with an N-terminal His-tag. For expression, transformed DH10B *E. coli* cells (Life Technologies) were grown in a shaking incubator (250 rpm) at 37°C in LB supplemented with 100 μg ml^−1^ ampicillin. When a cell density of OD_600_ ~0.4 was reached, cultures were either immediately induced with isopropyl-β-D-1-thiogalactopyranoside (IPTG, at final concentration of 1 mM), or transferred to a 25°C shaking incubator (250 rpm) and induced 30 min later with the same concentration of IPTG. After 5 hours of expression, cells were pelleted by centrifugation, and soluble and insoluble proteins were fractionated using the B-PER Bacterial Protein Extraction Reagent (Pierce) supplemented with protease inhibitor cocktail (Complete, EDTA-free, Roche), both according to the manufacturer’s protocol. Protein fractions were analyzed by Coomassie staining of SDS-PAGE gels and immunoblotting with Anti-PentaHis HRP Conjugate (Qiagen). Blots were stripped with Restore Western Blot Stripping Buffer (Pierce) before being re-probed with Streptavidin-HRP (Pierce). In all cases, apparent molecular weights were approximated by comparison with the Precision Plus Protein Dual-Color Standard (Bio-Rad).

### Protein purification

Soluble fractions from large-scale expression cultures (500 mL - 1 L) were prepared as described above and used for affinity purification of His-tagged proteins with Ni-NTA Agarose (Qiagen) according to the manufacturer’s protocol for native conditions. Following purification, proteins were buffer exchanged into PBS using PD-10 Desalting Columns (GE Healthcare Life Sciences) and quantified using the BCA Protein Assay Kit (Pierce).

### Circular permutation of miniSOG-Jα

A non-repetitive, codon-optimized (*E. coli*) DNA sequence encoding a tandem dimer of miniSOG-Jα was synthesized by PCR overlap-extension and used as a template for amplification of individual CPs. The resulting DNA fragments were inserted into a bacterial expression vector (pQE-80L, Qiagen) in-frame with an N-terminal His-tag. Expression was performed at 25°C as described above for miniSOG and miniSOG-Jα. After 5 hours of expression, cultures were pelleted by centrifugation, washed once with PBS, and resuspended in PBS at a normalized cell density of OD_600_=0.5. MiniSOG fluorescence was measured in a black 96-well plate using a fluorescence plate reader and imaged using a Maestro Imaging System (PerkinElmer). To confirm expression of individual CPs, total protein was extracted from normalized cell suspensions by sonication in PBS followed by addition of SDS to a final concentration of 1% (w/v) and heating (~50°C for 5 min) for solubilization of total protein. Insoluble cell debris was removed by centrifugation and the supernatant subsequently heated at 95°C for 5 min in 1X reducing SDS-PAGE loading buffer and analyzed by SDS-PAGE and Coomassie staining.

### Complementation assays in *E. coli*

DNA encoding mSOG_1-94_-NZip was inserted into pBAD/Myc-His A (Life Technologies), and DNAs encoding CZip-mSOG_95-106_ and CZip-mSOG-Jα_95-140_ were inserted into pBAD-18cm (Addgene). For protein co-expression, plasmids were sequentially transformed into DH10B *E. coli* and double-transformants were maintained using ampicillin (100 μg ml^−1^) and chloramphenicol (35 μg ml^−1^). Proteins were expressed at 25°C as described above for miniSOG and miniSOG-Jα, except using L-arabinose as the inducer (at a final concentration of 1% w/v) in place of IPTG. The soluble fraction of cell lysates was analyzed by Coomassie-staining of SDS-PAGE gels or used for affinity purification of Histagged complexes, both as described above.

### Complementation assays in mammalian cells

DNA sequences encoding reporter-tagged proteins were inserted downstream of a CMV promoter in a mammalian expression vector (pcDNA3.1, Life Technologies) that was modified to include an encephalomyocarditis virus (ECMV) IRES sequence driving expression of a fluorescent protein (mCherry, Citrine, or their mitochondria (mt)-targeted derivatives). Mammalian cells (HEK-293 or HeLa, as indicated in the text and figure captions) were cultured in imaging dishes with a coverslip bottom (P35G-0-14-C, MatTek Corporation) and transfected with DNA using Lipofectamine 3000 (Life Technologies) according to the manufacturer’s protocol. Cells were imaged by epifluorescence 24 - 48 hours following transfection in HBSS. For FKBP/FRB expressing cells, 100 nM rapamycin or DMSO was added to cultures ~16 hours prior to imaging. Imaging miniSOG in cells that also expressed Citrine required off-peak excitation for selective detection of each chromophore using the following optical settings: EX405/20, DM505, EM535/25.

### Imaging of α-syn interaction in cultured neurons

Cortical neurons were dissociated by papain from postnatal day 2 (P2) Sprague Dawley rats, and co-transfected with a total of 5.0 μg DNA (2.5 μg of each vector used) by electroporation using an Amaxa Nucleofection Device (Lonza). mSOG_1-94_ and mSOG-Jα_95-140_ were fused to the N-terminus and C-terminus of α-syn. Co-transfections were done with N- and C-terminus fusions, or C- and C-terminus fusions. The transfected neurons were plated on imaging dishes (P35G-0-14-C, MatTek Corporation) that were coated on the same day with poly-D-lysine. Neurons were cultured in Neurobasal A medium containing 1X B27 Supplements (both from Life Technologies), 2 mM GlutaMAX (Life Technologies), 20 U/mL penicillin, and 50 μg/mL streptomycin for 14 days prior to imaging. Neurons were imaged in HBSS containing 1X B27 Supplements, 25 mM glucose, 1 mM pyruvate, and 20 mM HEPES. All animal procedures were approved by the Institutional Animal Care and Use Committee of UC San Diego.

Confocal immunofluorescence images (1024 × 1024 pixels) were acquired on the Olympus Fluoview 1000 laser scanning confocal microscope using a 60X oil immersion objective with numerical aperture 1.42.

### *In vitro* analysis of split-miniSOG reversibility

A 17 amino acid TEV-protease cleavable linker (GSGSGENLYFQSGSGSG) was inserted into miniSOG and miniSOG-Jα between Gly94 and Glu95, and the resulting proteins were fused with a SUMO tag at their C-termini, generating TC-miniSOG and TC-miniSOG-Jα, respectively. To generate TC-Venus, a 20 amino acid cleavable linker (GSGSGSGENLYFQSGSGSGG) was inserted into the fluorescent protein Venus between residues Gln158 and Gln159. Proteins were expressed at 25°C from an arabinose inducible promoter as described above, except expression was done overnight. The soluble fractions of cell lysates were used for affinity purification of His-tagged proteins as described above. Proteins were cleaved using ProTEV Plus protease (Promega) following the manufacturer’s protocol with modifications: 2.5-times the recommended amount of substrate protein (40 μg per 100 μl reaction) and 1.5-times the amount to recommended enzyme (1.5 μl per 100 μl reaction) was used in to cleave proteins in 200 μl reaction volumes. Cleaved and uncleaved samples were analyzed using a fluorescence plate reader in a black 96-well plate. An aliquot of each sample was collected prior to fluorescence measurement and analyzed by SDS-PAGE followed by Coomassie staining.

### Confocal fluorescence imaging and photo-oxidation

Mammalian cells were cultured in dishes with a coverslip glass bottom (P35G-0-14-C, MatTek Corporation) and transfected with DNA using Lipofectamine 3000 as described above. Proteins were allowed to express for 24 – 48 hours before cells were rinsed with pre-warmed HBSS and fixed using pre-warmed 2% (w/v) glutaraldehyde (Electron Microscopy Sciences) in 0.1 M sodium cacodylate buffer, pH 7.4 (Ted Pella Incorporated) for 5 minutes at 37 °C and then on ice for 1 hour. Subsequently, cells were rinsed on ice 3-5 times using chilled cacodylate buffer and treated for 30 minutes on ice in a blocking solution (50 mM glycine, 10 mM KCN, and 5 mM aminotriazole in 0.1 M sodium cacodylate buffer, pH 7.4) to reduce nonspecific background precipitation of DAB. Cells were imaged and photooxidized using a Leica SPE II inverted confocal microscope outfitted with a stage chilled to 4 °C. Confocal fluorescence and transmitted light images were taken with minimum exposure to identify transfected cells for correlative light microscopic imaging. For photooxidation, DAB (3-3’-diaminobenzidine tetrahydrochloride, Sigma-Aldrich) was dissolved in 0.1 N HCl at a concentration of 5.4 mg ml^−1^ and subsequently diluted ten-fold into sodium cacodylate buffer (pH 7.4, with a final buffer concentration of 0.1 M), mixed, and passed through a 0.22 μm syringe filter before use. DAB solutions were freshly prepared on the day of photooxidation and placed on ice and protected from light before being added to cells. Regions of interest were identified by fluorescence and images were recorded with care to avoid sample photo-bleaching. The samples were then illuminated through a standard FITC filter set (EX470/40, DM510, BA520) with intense light from a 150 W xenon lamp. Illumination was stopped as soon as an optically-dense reaction product began to appear in place of the green fluorescence, as monitored by transmitted light (typically 3-8 min, depending on the initial fluorescence intensity, the brightness of the illumination, and the optics used).

### Electron Microscopy

Multiple areas on a single dish were photooxidized as described in the preceding section. Subsequently, cells were placed on a bed of ice and washed using ice-cold cacodylate buffer (5×2 minutes) to remove unpolymerized DAB. After washing, cells were post-fixed with 1% osmium tetroxide (Electron Microscopy Sciences) in 0.1 M sodium cacodylate buffer for 30 minutes on ice, then washed with ice-cold cacodylate buffer (3×2 minutes) and rinsed once in ice-cold distilled water. The samples were then dehydrated with an ice-cold graded ethanol series (20%, 50%, 70%, 90%, 100%, 100%, 3 minutes each) and washed once in room temperature anhydrous ethanol (3 minutes). Samples were then infiltrated with Durcupan ACM resin (Electron Microscopy Sciences) using a 1:1 solution of anhydrous ethanol:resin for 30 minutes on a platform with gentle rocking, then with 100% resin overnight with rocking. The next day, the resin was removed from dishes (by decanting and gentle scraping with care to avoid touching cells), replaced with freshly prepared resin (3×30 minutes with rocking), and polymerized in a vacuum oven at 60°C for 48 hours. Subsequently, photooxidized areas of interest were identified by transmitted light, sawed out using a jeweler’s saw, and mounted on dummy acrylic blocks with cyanoacrylic adhesive. The coverslip was carefully removed and ultrathin sections (80 nm thick) were cut using a diamond knife (Diatome). Electron micrographs were recorded using a JEOL 1200 EX transmission electron microscope operating at 60/80 kV. For electron tomography, thicker sections (200-250 nm) were imaged using an FEI Titan high base microscope operated at 300kV; micrographs were produced using a 4k x 4k Gatan CCD camera (US4000). Colloidal gold particles (5 and 10nm diameter) were deposited on each side of the sections to serve as fiducial markers. For each section, double-tilt series were collected using the SerialEM package. For each series, the sample was tilted from -60 to +60 degrees, every 0.5 degree. Tomograms were generated using an iterative reconstruction procedure^13^.

Energy Filtered Transmission Electron microscopy (EFTEM) was performed with a JEOL JEM-3200EF transmission electron microscope operating at 200 KV, equipped with an in-column Omega filter. The samples were pre-irradiated at a low magnification of 100X for about 30 minutes to stabilize the sample and minimize contamination. The elemental maps were obtained at the M_4,5_ core-loss edge, the onset of which occurs at 883 cerium (Ce). The EFTEM images of the pre and post-edges were obtained using the 3-window method and a slit width of 40 eV. The pre-edges were obtained at 790 and 850 eV, and the post-edge was obtained at 908 eV. The elemental maps were computed using the EFTEM-TomoJ plug-in of ImageJ ^21^, using the power law fit.

The EFTEM images were acquired using the direct detection device DE-12 from Direct Electron (San Diego, CA, USA). Each of the pre-edge and post-edge image, is the summed image of 14 and 11 individual drift corrected and aligned images, respectively for the two sets of data shown in the figure 3. This technique mitigates effects of sample drift and other microscope instabilities over time, and the details of such an image acquisition and processing is described elsewhere^13-22^. The individual images were acquired for an exposure of 125 s for a frame rate of 0.04 frames/sec.

## Acknowledgments

In memory of Roger Y. Tsien (RYT) who inspired and co-supervised this work with MHE. RYT passed away in August 2016. We thank H. Hakozaki and P. Steinbach for advice regarding optical microscopy, and P. Nguyen for assistance with sample preparation for electron microscopy. This work was supported by NIH grants to RYT, MHE, SRA and VLR (NS027177), to SRA/DB (R01 GM086197), to DB and MHE (Branfman Family Foundation), and to MHE (P41 GM103412) for support of the National Center for Microscopy and Imaging Research. RYT was an Investigator in the Howard Hughes Medical Institute.

## Author Contributions

J.T.N., D.B., S.P.L., J.H., Q.X., S.P., M.M., R.R., and V.L-R. performed experiments. J.T.N., D.B., S.R.A., and M.H.E. wrote the paper. All authors analyzed data and edited the manuscript.

## Competing Financial Interests

The authors declare no competing financial interests.

## Supplementary Material

**Supplementary Figure 1.**
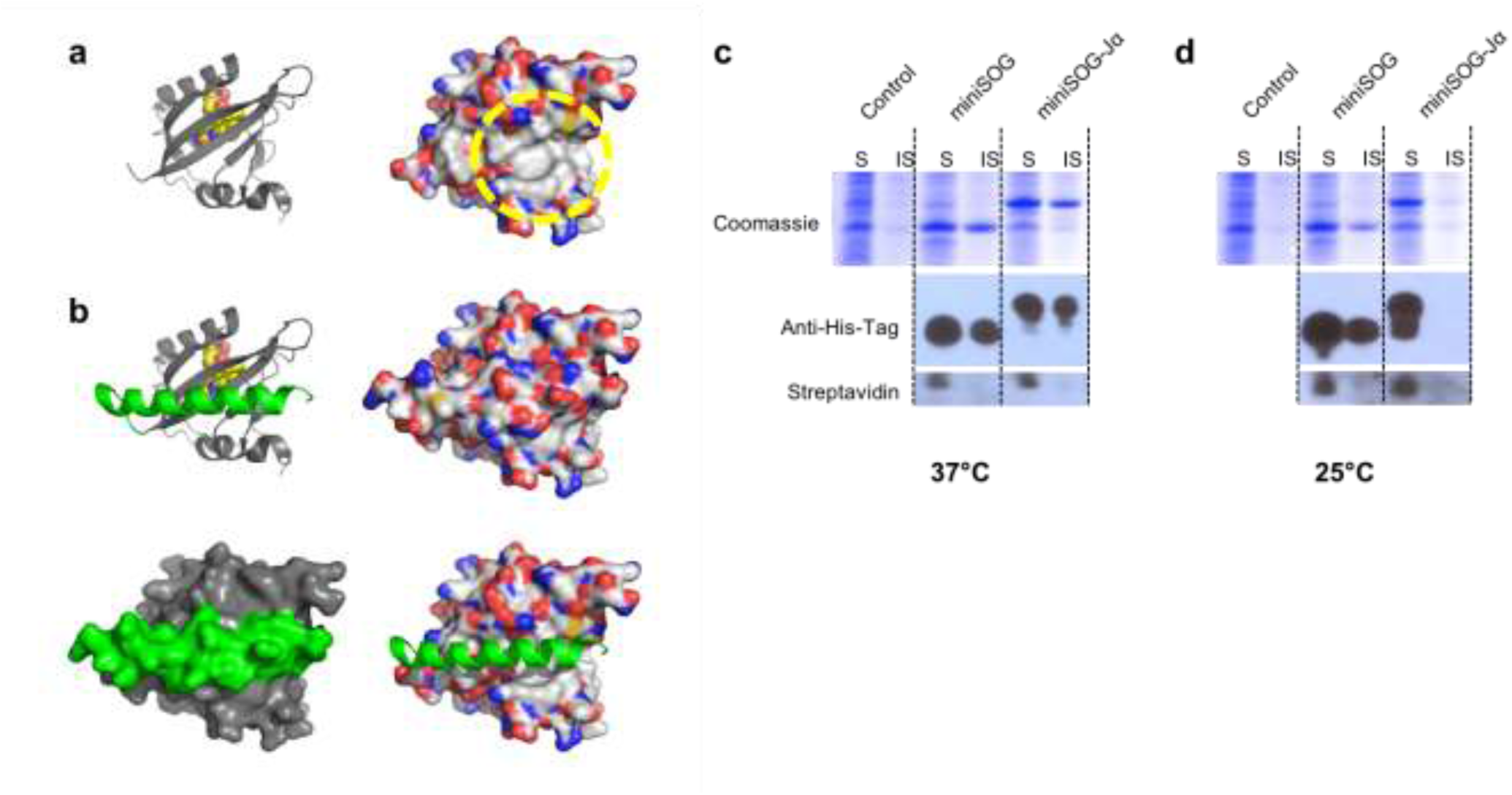
| Inclusion of the Jα-helix increases the solubility of miniSOG when rapidly and highly overexpressed in *E. coli*. (**a**) Structures of the LOV2 domain from oat (PDB: 2V0U) shown without Jα-helix residues. A small, solvent-exposed hydrophobic patch is highlighted (yellow dotted line, right image). Cartoon images are displayed with FMN shown as spheres (left image). Coloring by atom (right image): C is light gray, O is red, N is blue, and S is yellow. (**b**) Structures of the same domain displayed with the Jα-helix shown (green), which sequesters the surface-exposed hydrophobic patch. (**c-d**) Comparison of inclusion body formation between miniSOG and miniSOG-Jα. Briefly, His-tagged versions of the proteins were over-expressed in *E. coli* from an IPTG-inducible promoter at either 37° or 25°C. Expression at both temperatures led to high levels of recombinant protein (~20% of total cellular protein content). Soluble and insoluble cell fractions were subsequently prepared and analyzed by Coomassie staining of SDS-PAGE gels as well as by Western blotting with an anti-His-tag antibody. In cells that were induced at 25°C, miniSOG-Jα was completely soluble and was not detected in the insoluble fraction. To assess the quality of the cell fractionation procedure, Western membranes were stripped and re-probed with streptavidin-HRP in order to reveal biotin-carboxyl carrier protein (~17 kDa), a soluble and endogenously biotinylated *E. coli* protein. Streptavidin-HRP signal was detected only in soluble fractions, indicating a successful separation of soluble and insoluble proteins.

**Supplementary Figure 2.**
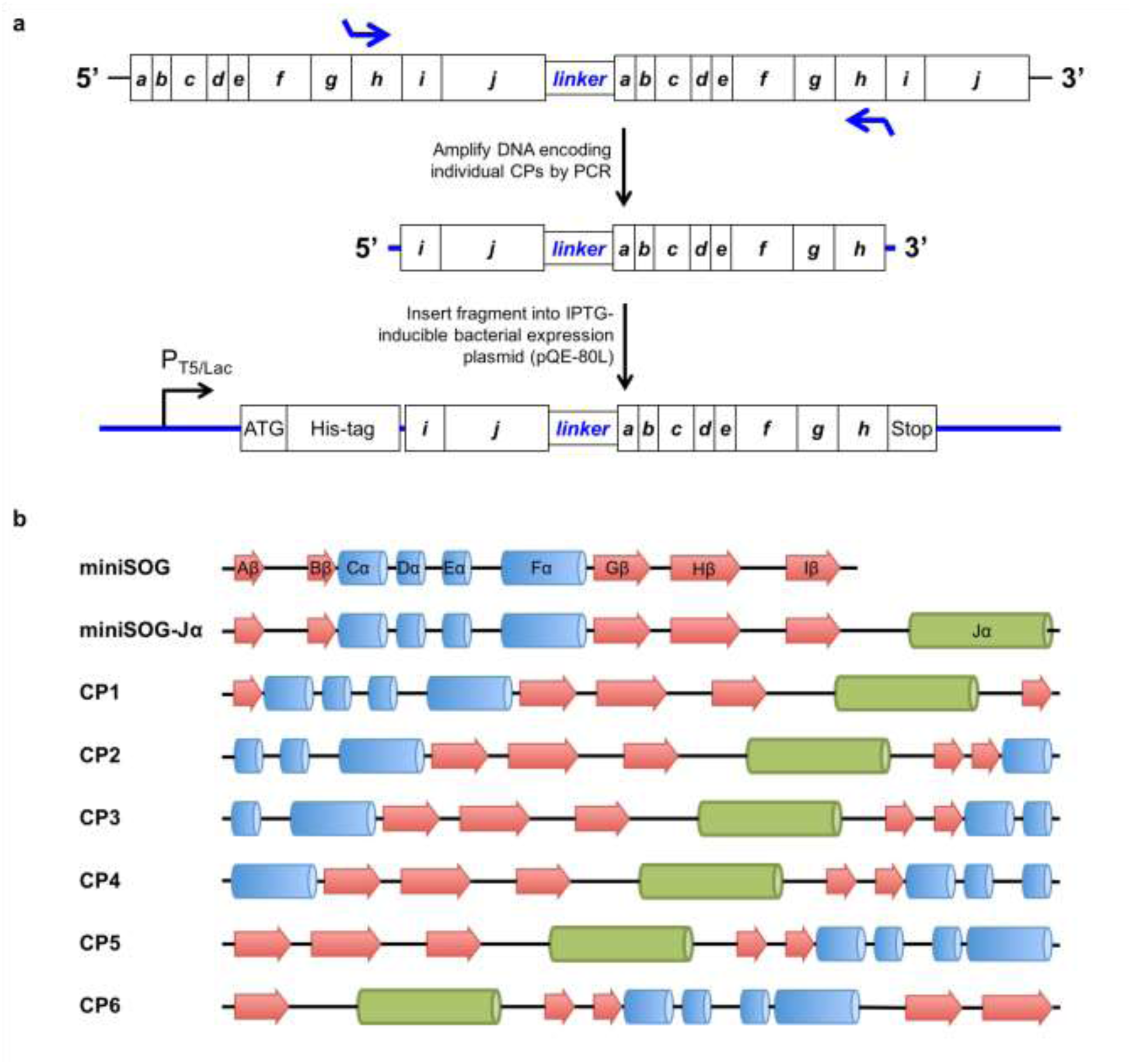
| Generation of miniSOG-Jα CP pool. (**a**) Schematic of the cloning strategy used to generate DNA sequences encoding the individual CPs. A non-repetitive DNA sequence encoding a miniSOG-Jα tandem dimer was used as a template from which monomeric CP genes were amplified using PCR. Generation of the CP6 gene is depicted. (**b**) Secondary structural elements of miniSOG, miniSOG-Jα, and CP1 - CP6. Beta sheets are shown as arrows, and helices are shown as cylinders. The Jα-helix is shown as a green cylinder.

**Supplementary Figure 3.**
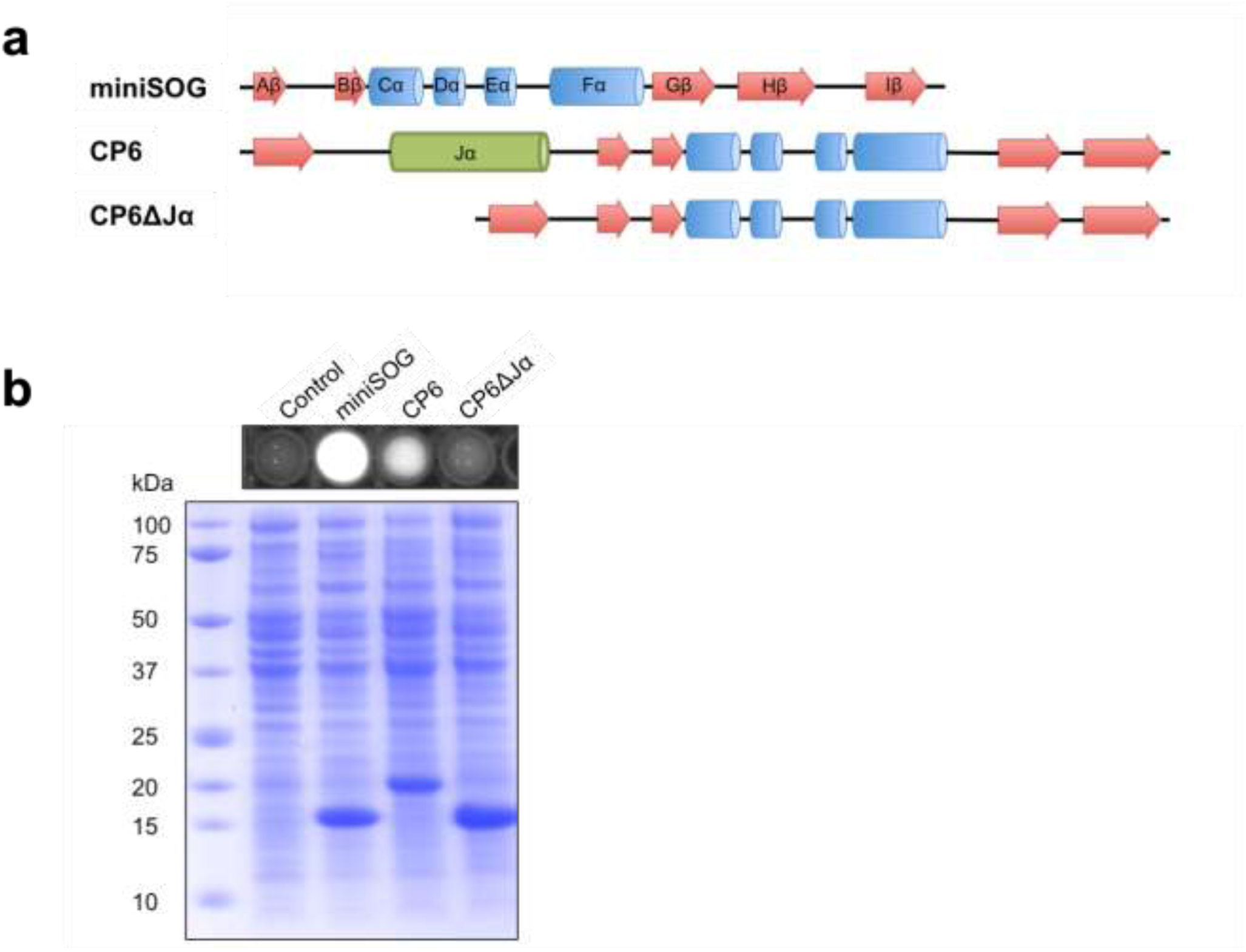
| The Jα-helix aids in the proper folding of CP6. (**a**) Schematic representation of the secondary structural elements of miniSOG, CP6, and CP6ΔJα. (**b**) Fluorescence detection of *E. coli* cells expressing the indicated proteins (top) and Coomassie-stained SDS-PAGE gel of corresponding total protein extracts, including both the soluble and insoluble forms of each species (bottom). Calculated protein masses of the His-tagged proteins: miniSOG = 15.23 kDa, CP6 = 19.11 kDa, CP6ΔJα = 15.34 kDa.

**Supplementary Figure 4.**
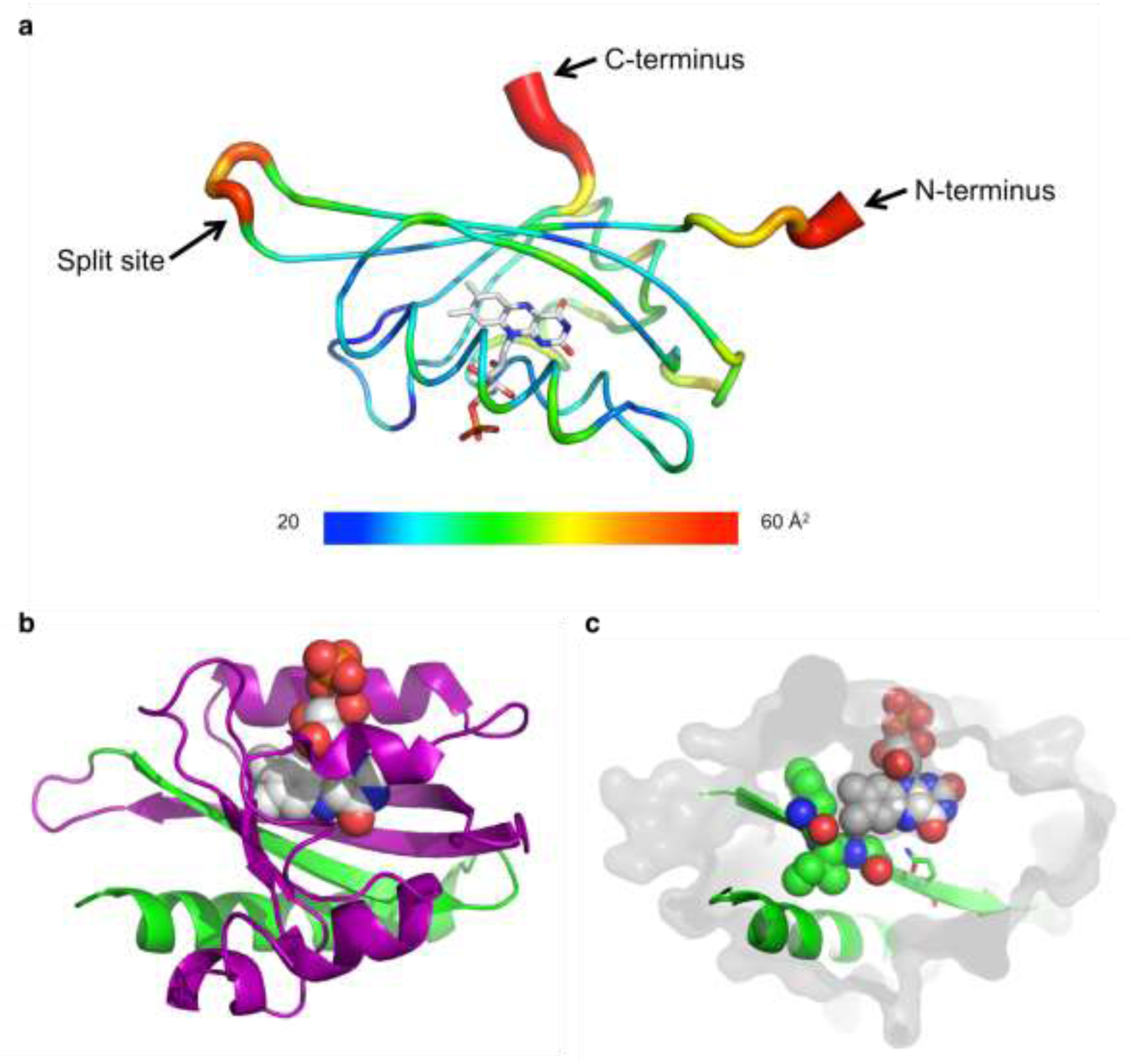
| Structural analyses of miniSOG/miniSOG-Jα bisection. (**a**) Wire diagram on the LOV2 domain (PDB: 4EEP) with relative B-factor values represented as a heat map (blue to red: 20 - 60 Å^2^) and also by wire diameter. B-factor values correspond to the displacement of atomic positions from the mean per atom, as calculated from crystallographic x-ray diffraction data. High B-values are associated with regions of high flexibility. FMN is shown as sticks. Arrows indicate the N-terminus, C-terminus, and the identified split-site (between Gly94 and Glu95). The image was generated using Chimera (https://www.cgl.ucsf.edu/chimera/). (**b**) Structure of the LOV2 domain (PDB: 2V0U) with residues corresponding to mSOG_1-94_ shown in purple, and residues corresponding to mSOG-Jα_95-140_ shown in green. FMN (spheres) makes direct contact with regions of both fragments. (**c**) While mSOG_1-94_ forms the bulk of the FMN binding-pocket (gray surface), residues on mSOG_95-106_/ mSOG-Jα_95-140_ (green) make contact with the cofactor through van der Waals interactions (residues 99 – 101, spheres) and through hydrogen bonding between the side-chain N of Glni03 (shown in sticks) and O_4_ of the FMN isoalloxazine ring. mSOG_95-106_ is a substructure of mSOG-Jα_95-140_ and lacks residues corresponding to the Jα-helix.

**Supplementary Figure 5.**
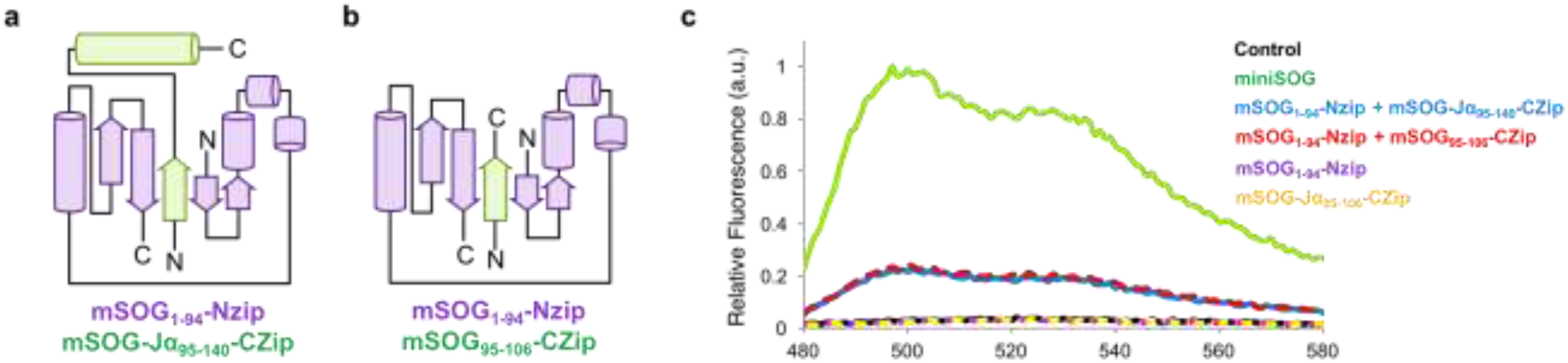
| Comparison of mSOG_1-94_JmSOG-Jα_95.140_ and mSOG_1-94_ and mSOG-_95-106_ in *E. coli* and mammalian cells. (**a**) Topology of mSOG_1-94_ (purple) and mSOG-Jα_95-140_ (green) tagged with leucine zipper sequences (Nzip and Czip). (**b**) Topology of mSOG_1-94_ (purple) and mSOG_95-106_ (green) tagged with Nzip and Czip. (**c**) Expression of individual miniSOG fragments in *E. coli* is insufficient for upregulation of cellular FMN biosynthesis. Fluorescence emission spectra from *E. coli* cells expressing leucine zipper sequences (Nzip and Czip) tagged with miniSOG/miniSOG-Jα fragments. Independent expression of mSOG_1-94_-Nzip, mSOG-Jα_95-140_-Czip did not result in detectable fluorescence above background. Co-expression of mSOG_1-94_-Nzip and mSOG-Jα_95-140_-Czip, or mSOG_1-94_-Nzip and mSOG_95-106_-Czip resulted in comparable levels of fluorescence with spectral profiles resembling that of miniSOG.

**Supplementary Figure 6.**
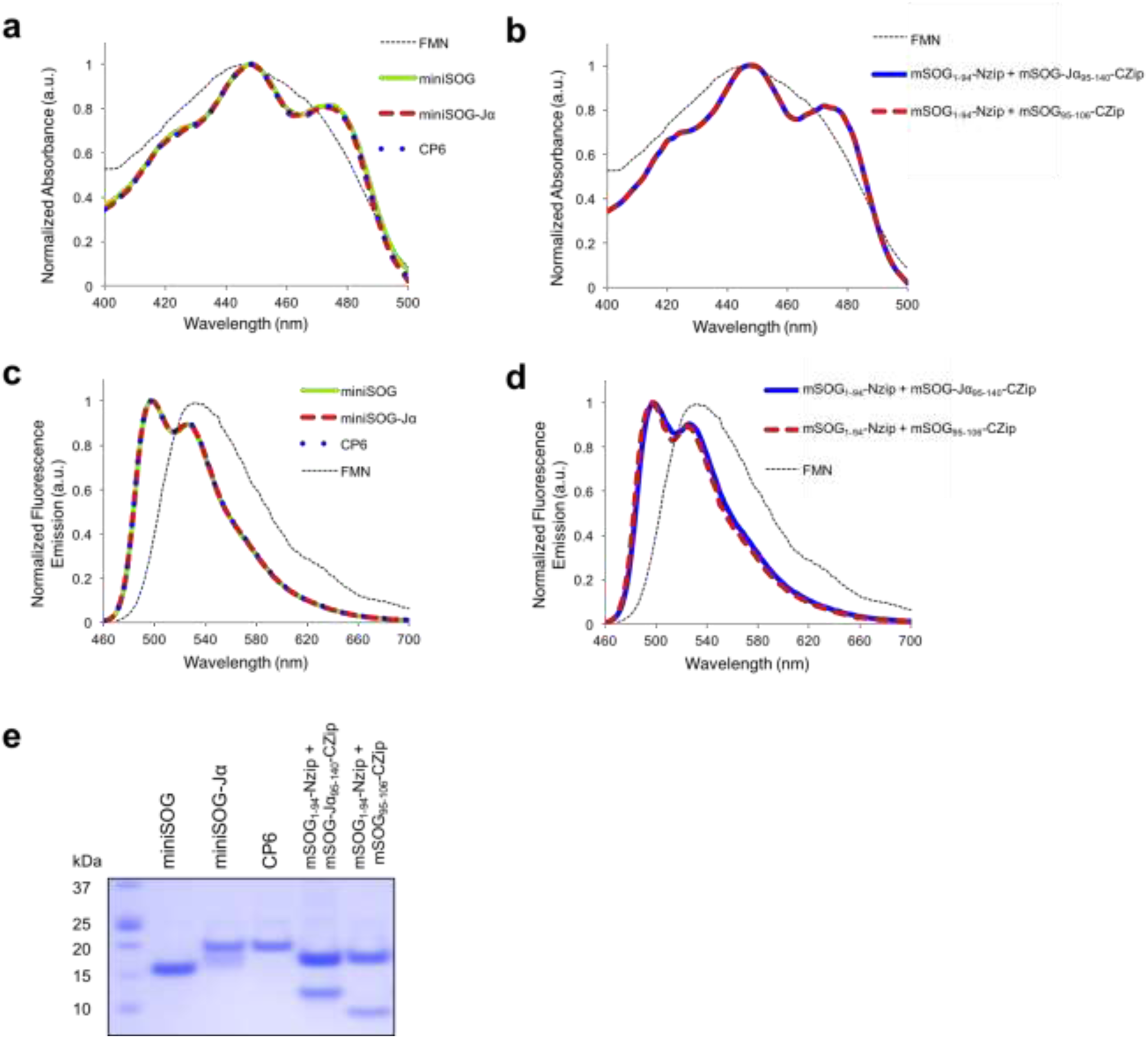
| FMN remains bound to miniSOG-Jα, CP6, and reconstituted complexes following purification. Normalized absorption and fluorescence emission spectra of free FMN and purified miniSOG, miniSOG-Jα, CP6 (**a, c**), as well as mSOG_1-94_-Nzip/mSOG-Jα_95-140_-Czip and mSOG_1-94_-Nzip/mSOG_-95-106-_Czip complexes (**b, d**). LOV-domain bound FMN exhibits characteristic vibronic structure (multiple peaks) that is not observed with the free chromophore. (**e**) Coomassie-stained SDS-PAGE gel of the purified proteins used to collect the spectra shown in (a-d). Calculated protein masses: miniSOG (with myc and His-tag) = 15.23 kDa, miniSOG-Jα (with His-tag) = 19.11 kDa, CP6 (with His-tag) = 19.11 kDa, mSOG_1-94_-Nzip (with His-tag) = 15.98 kDa, mSOG-Jα_95-140_-Czip (untagged) = 10.66 kDa, mSOG_-95-106-_Czip (untagged) = 6.90 kDa.

**Supplementary Figure 7.**
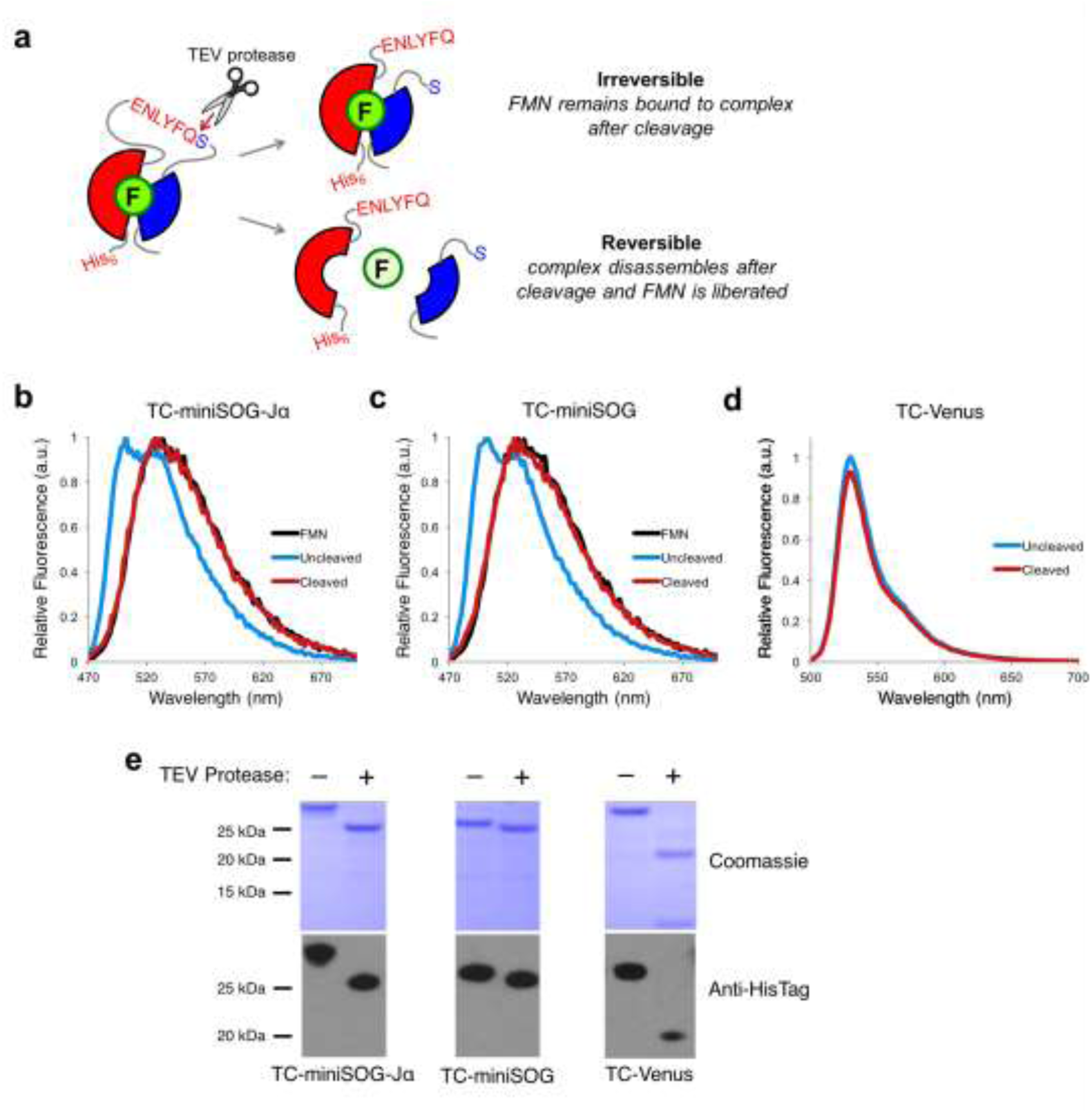
| FMN is liberated from reconstituted miniSOG/miniSOG-Jα complexes in the absence of a stabilizing template. (**a**) Schematic diagramming TEV-protease cleavage of full-length test proteins. A cleavable sequence was inserted at the split site into full-length miniSOG-Jα, miniSOG, and Venus, generating the TEV-protease cleavable (TC) proteins TC-miniSOG-Jα, TC-miniSOG, and TC-Venus. (**b-c**) Normalized emission spectra are shown for uncleaved (blue lines), TEV-protease cleaved TC-miniSOG-Jα and TC-miniSOG (red lines), and free FMN alone (black lines). Uncleaved TC-miniSOG-Jα and TC-miniSOG samples exhibited emission spectra corresponding to LOV domain-bound chromophore (blue traces), indicating that FMN was associated with the protein prior to cleavage. Following digestion with TEV protease, the samples exhibited emission spectra corresponding to liberated FMN (red traces) matching emission of free FMN in solution (black traces), thus indicating that FMN is liberated from protein complex in the absence of a stabilizing template. (**d**) The control protein TC-Venus is similarly fluorescent before and after digestion with TEV-protease. Relative emission spectra from TC-Venus before and after cleavage are shown. (**e**) Complete cleavage of the test proteins was verified by Coomassie staining of SDS-PAGE gels and immunoblotting with detection for N-terminal His-tags.

**Supplementary Figure 8.**
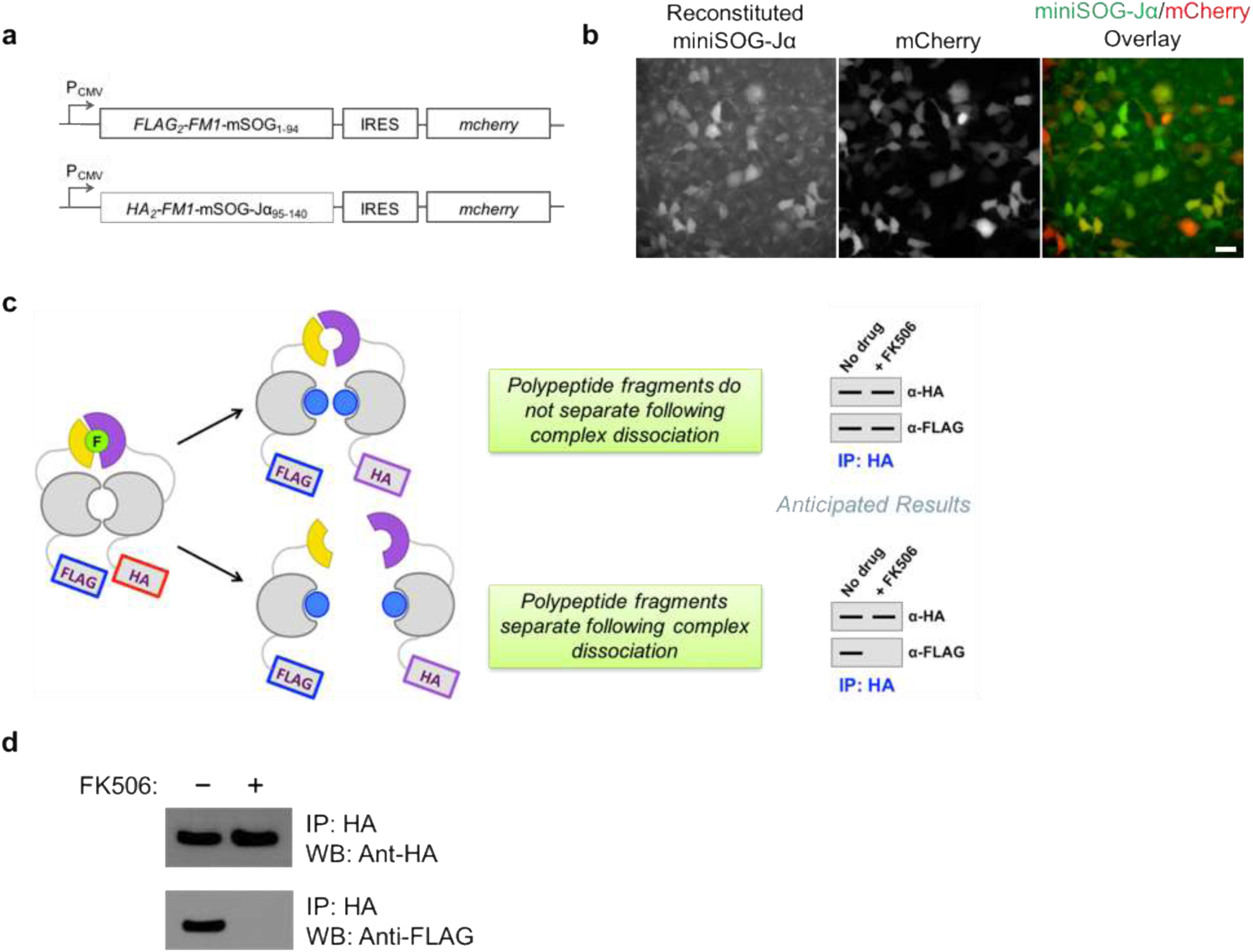
| miniSOG_1-94_ and miniSOG-Jα_95-140_ physically dissociate from one another following unbinding of fused interaction partners. (**a**) Schematic of the DNA constructs encoding tagged version of the self-associating FKBP14 mutant F_M1_ (FLAG_2_-F_M1_-mSOG_1-94_ and HA_2_-F_M1_-mSOG-Jα_95-140_, respectively). (**b**) Co-transfection of HEK293 with the constructs resulted in detectable fluorescence from reconstituted miniSOG-Jα, indicating that F_M1_ self-association is sufficient to drive domain reassembly. Transfected cells were identified by detection of mCherry. Scale bar, 30 microns. (**c**) Schematic diagramming the anticipated results following immunoprecipitation (IP) analysis. miniSOG_1-94_ and miniSOG-Jα_95-140_ are shown as yellow and purple shapes. F_M1_ spontaneously forms dimers (gray shapes in cartoon) in the absence of an FKBP ligand—thus, we anticipated that IP of HA_2_-F_M1_-mSOG-Jα_95-140_ (using anti-HA antibody) would result in co-IP of FLAG_2_-F_M1_-miniSOG_1-94_. In addition, we anticipated that co-IP could be abolished by FK506 (blue dots, a ligand that binds and dissociates FM1 dimers) if the interaction between the fragments was reversible. (**d**) Immunoblot detection of HA_2_-F_M1_-mSOG-Jα_95-140_ and FLAG_2_-F_M1_-mSOG_1-94_ in the elution fractions of anti-HA IPs done in the presence and absence of FK506. FLAG_2_-F_M1_-mSOG_1-94_ did not co-IP with HA_2_-F_M1_-mSOG-Jα_95-140_ in the presence of FK506, indicating that the reconstituted miniSOG-Jα does not physically bind F_M1_ proteins together following their drug-induced dissociation.

**Supplementary Figure 9.**
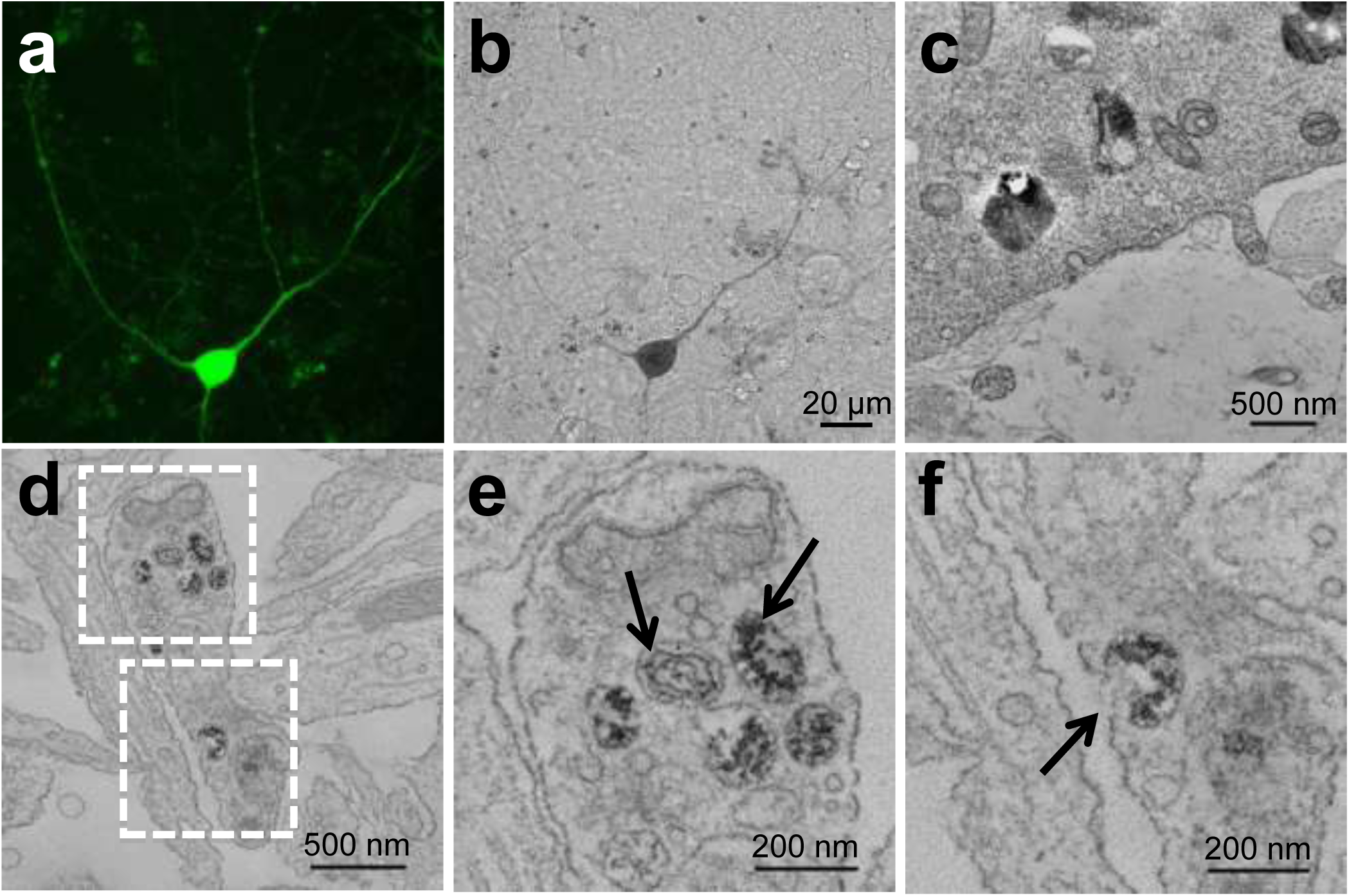
| Alpha-synuclein aggregates in the cell body and dendrites of neurons expressing the reconstituted miniSOG-Jα. (**a**) Confocal fluorescence image (maximum intensity projection) of live cultured neurons co-expressing the tagged α-syn proteins. A diffuse miniSOG fluorescent signal is observed throughout the neuron. Corresponding transmitted light image (**b**) was acquired post-fixation after 3 minutes and 30 seconds illumination in the presence of DAB. (**c-f**) TEM images of α-syn aggregates in the neuronal cell body (**c**) and dendrite (**d**). EM micrographs in (**e**) and (**f**) corresponds to white boxes in (**d**). Black arrows points at α-syn aggregates contained in membrane-limited organelles, fusing to the plasma membrane suggesting release to the extracellular compartment.

**Supplementary Figure 10.**
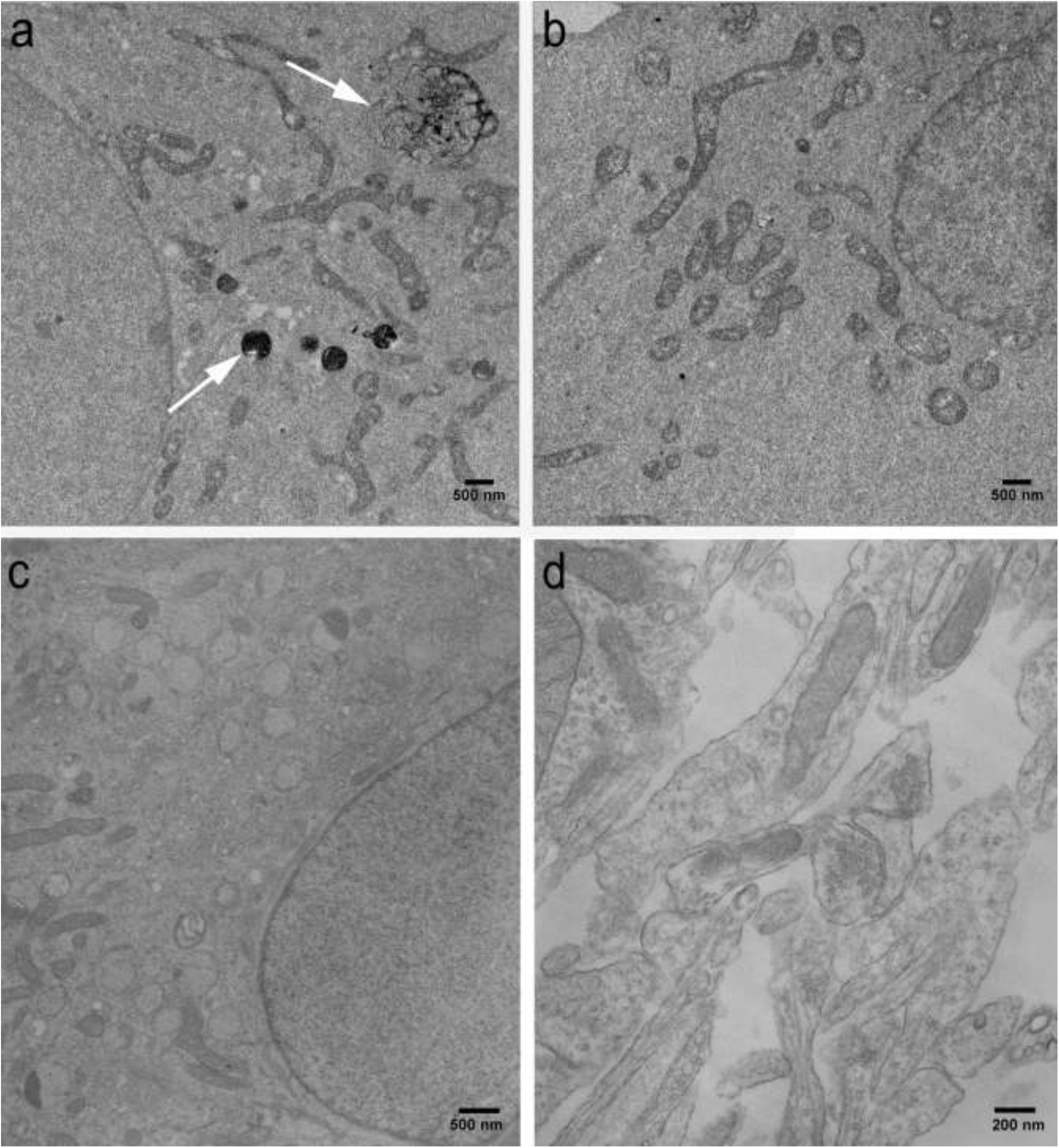
| Comparison of photooxidized and non-photooxidized neurons expressing tagged α-syn proteins, and photooxidized untransfected neurons. EM micrographs of cell bodies of photooxidized neurons co-expressing mSOG_1-94_ and mSOG-Jα_95-140_ fusions of α-syn (**a**), non-photooxidized neurons co-expressing the same proteins (**b**), and photooxidized untransfected neurons (**c** and **d**). Stained α-syn aggregates (white arrows) are observed only in the photooxidized expressing neurons.

**Supplementary Figure 11.**
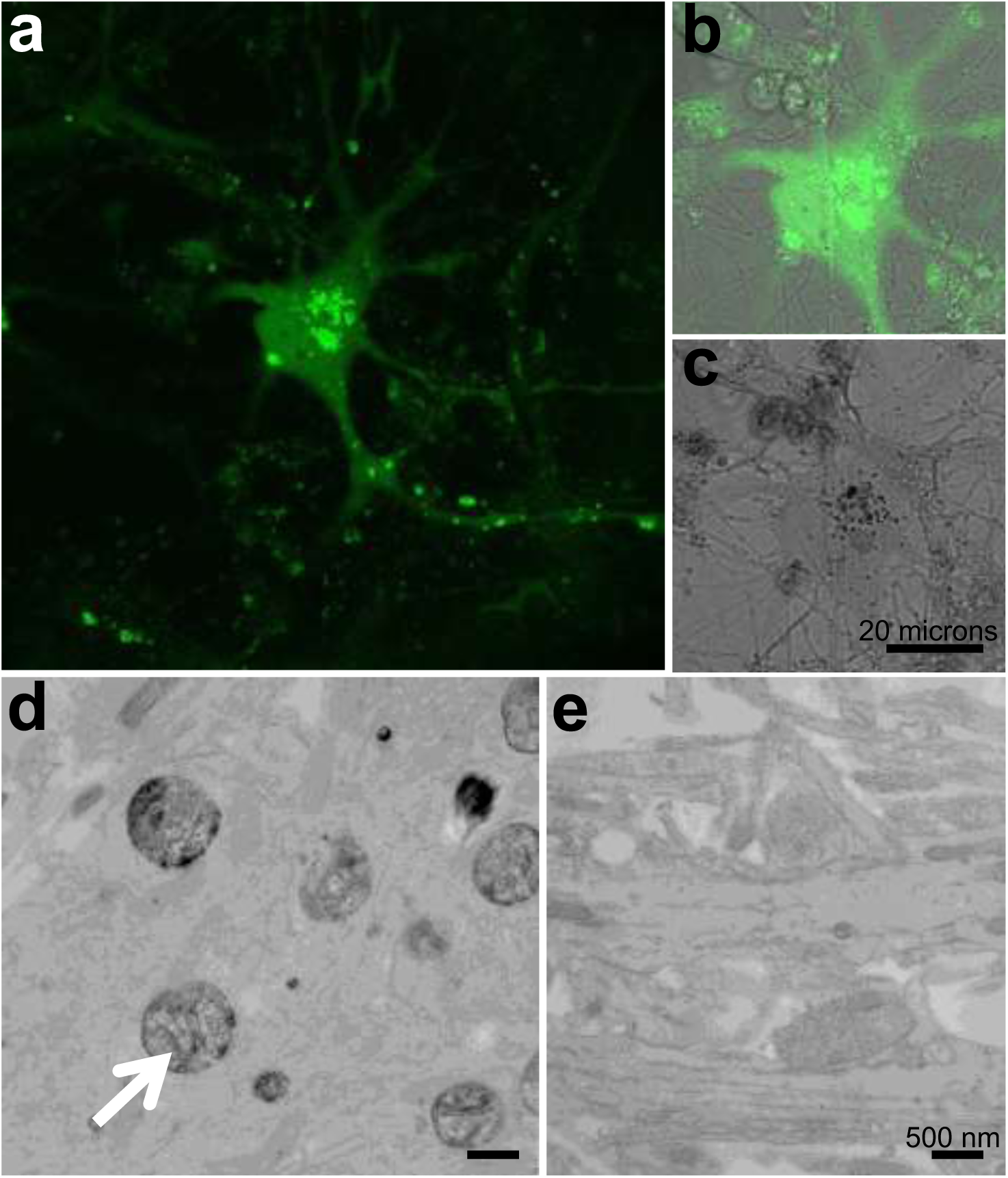
| Oligomeric A53T familial mutant displays distribution characteristics similar to wild type α-syn. (**a**) Confocal fluorescence image of live cultured neurons co-expressing the A53T tagged proteins, (**b**) Fluorescence image overlaid on the transmitted light image before photooxidation of DAB. (**c**) Following photooxidation, the appearance of the optically visible DAB reaction product correlates with the detected reconstituted miniSOG fluorescence in (**a, b**). (**d**) Electron micrograph showing labeled α-syn fibrillar structures (white arrow) similar to the ones observed for wild type α-syn. (**e**) No labeling was observed at presynaptic terminals where α-syn aggregates were not detected.

**Supplementary Figure 12.**
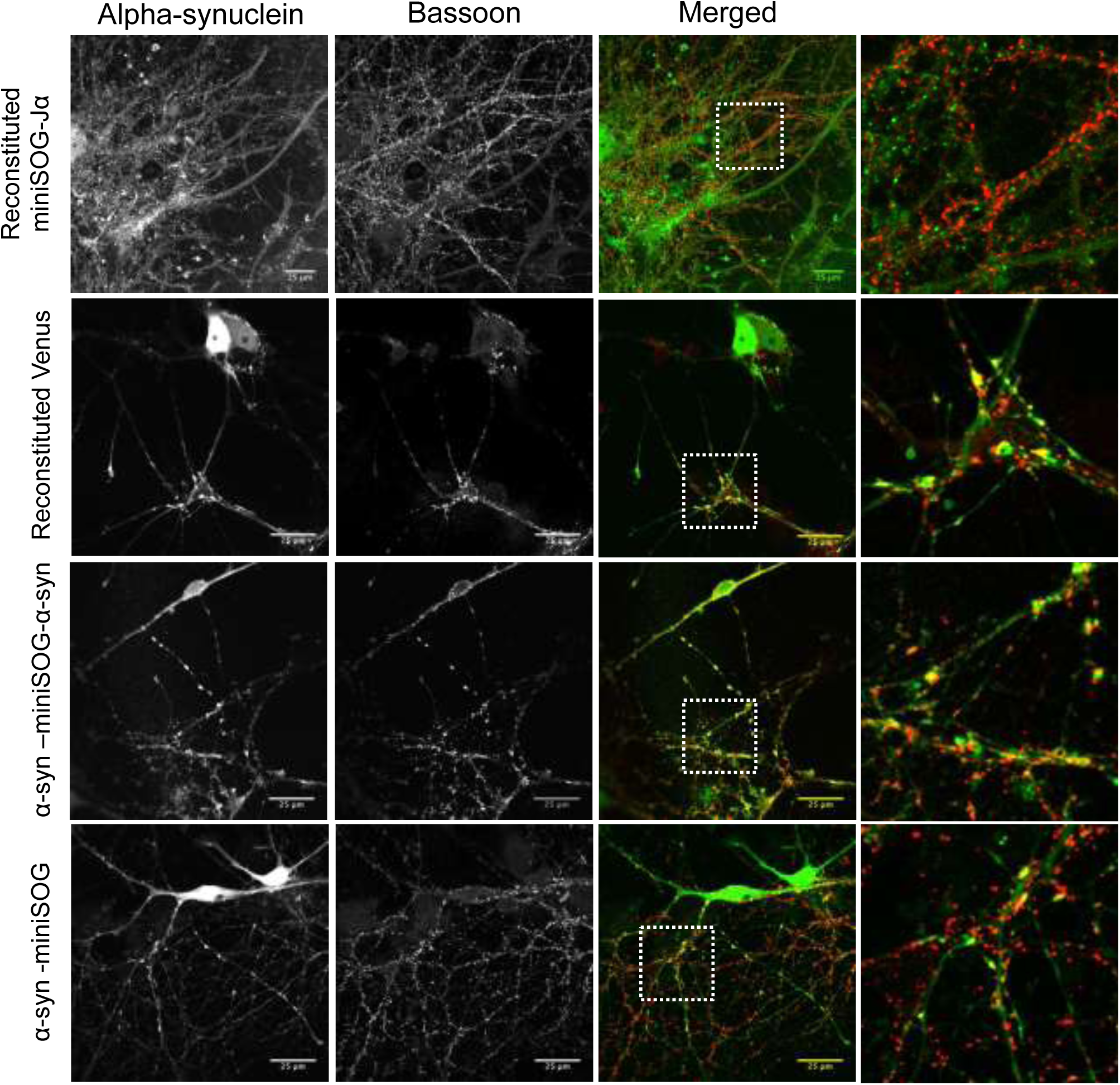
| Oligomeric wild-type α-syn is excluded from presynaptic terminals. Confocal images (single confocal planes) showing the localization of the α-syn proteins fused to different tags (first column, intrinsic fluorescence), the labeling for Bassoon (presynaptic marker, second column), the merged images (third column) and an enlargement of the boxed areas shown in the white squares (last column to the right). Comparison of the reconstituted miniSOG-Jα fluorescence with reconstituted Venus reveals no localization of the reconstituted miniSOG-Jα to presynaptic terminals opposite to the reconstituted Venus. Fusion of two α-syn proteins to the N- and C-termini of miniSOG respectively (α-syn-miniSOG-α-syn) mimics the irreversible bonding of the reconstituted Venus and shows strong localization to presynaptic terminals. Single α-syn-miniSOG fusion proteins are observed at presynaptic terminals.

**Supplementary Figure 13.**
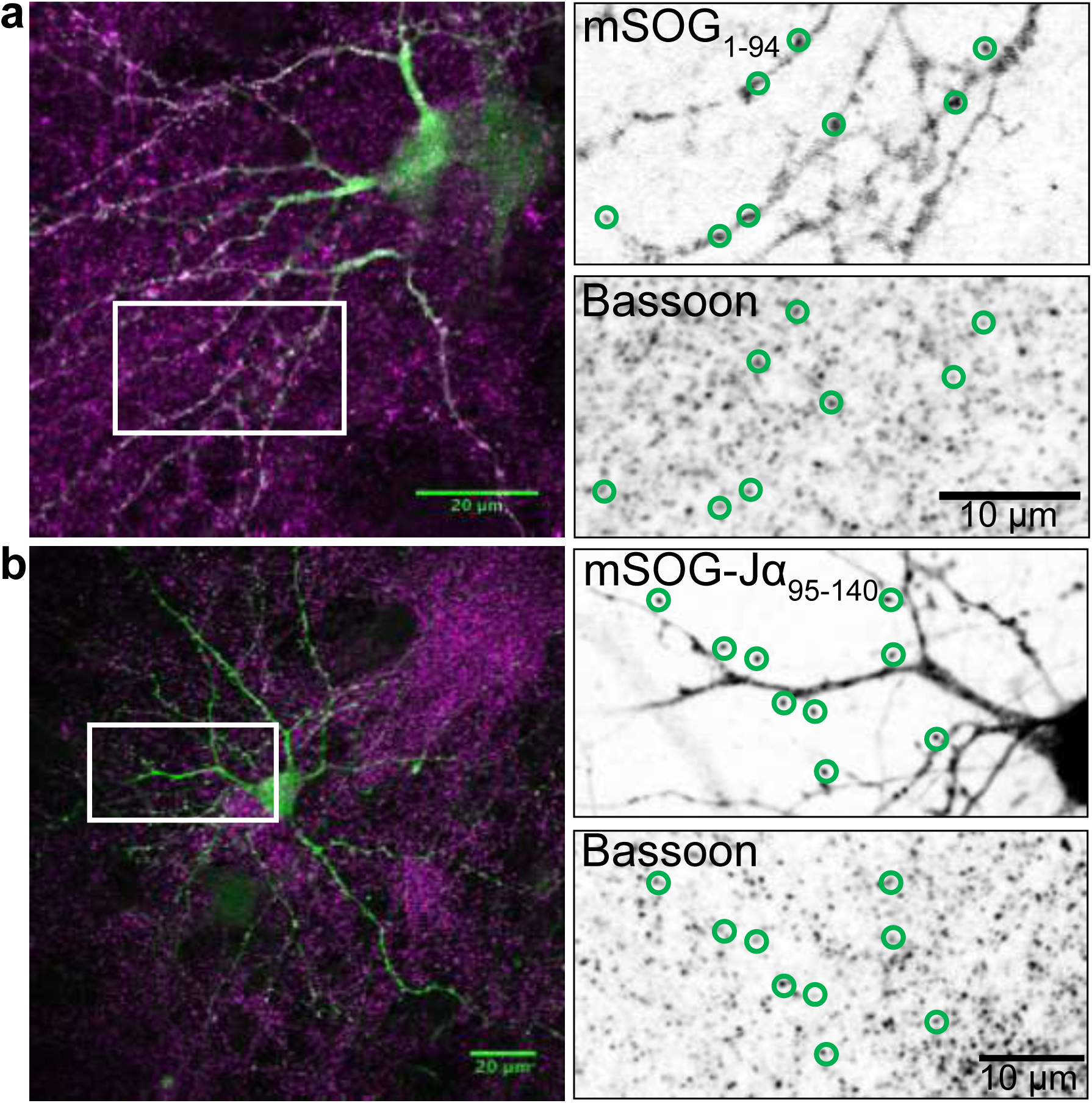
| Individual α-syn proteins fused to each split-miniSOG-Jα-tagged fragment are localized to presynaptic terminals. (**a**) Confocal images of rat cortical neurons coexpressing α-syn fused to mSOG_1-94_ (purple) and mSOG-Jα_95-140_. Immunolabeling of mSOG_1-94_ using an antibody that recognizes this fragment (RRX secondary antibody detection, shown in green in the merged image) and Bassoon (presynaptic marker, AX647 secondary antibody detection, shown in magenta in the merged image) demonstrate that the single-fragment fusions did not disrupt the ability of the α-syn proteins to traffic all the way to the presynaptic terminals. In the single channel confocal images (right column), the color table has been inverted for better visualization of the individual fluorescence channels. Green circles highlight corresponding labeling of α-syn proteins fused to each fragment colocalizing at presynaptic terminals. (**b**) Similarly, immunolabeling of mSOG-Jα_95-140_ was done using an antibody to an additional epitope, HA, introduced in the fusion protein (RRX secondary antibody detection, shown in green in the merged image).

**Supplementary Figure 14.**
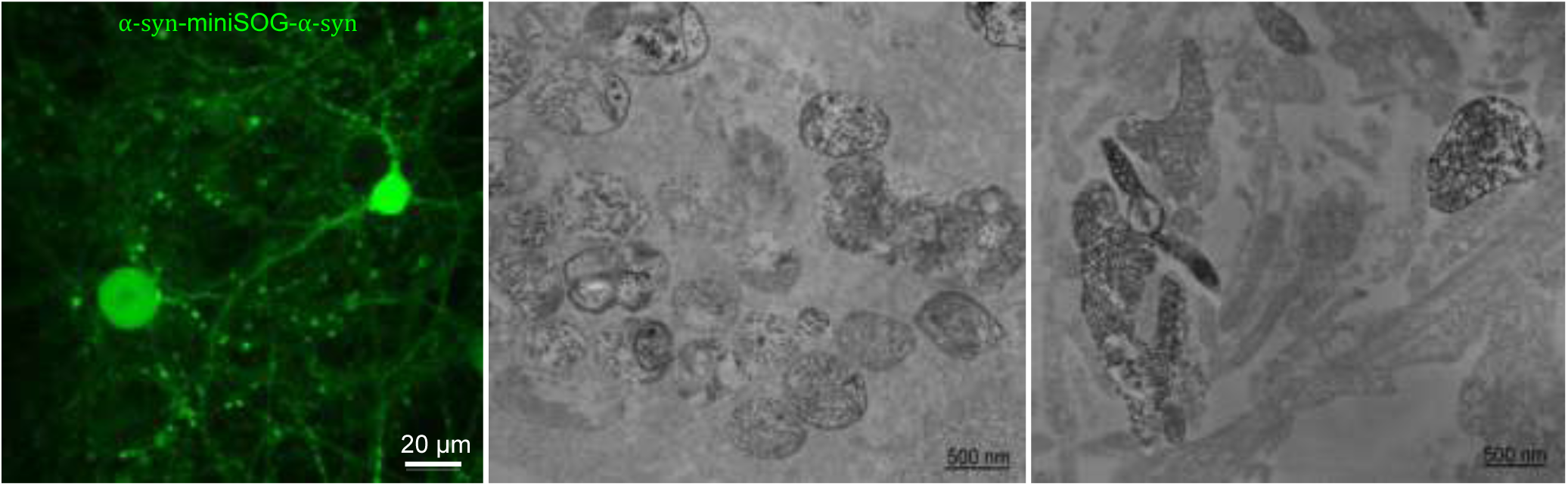
| Fusion of two α-syn proteins to the N- and C-termini of miniSOG respectively (α-syn-miniSOG-α-syn) mimics the irreversible bonding of the reconstituted Venus and shows strong labeling in presynaptic terminals. Confocal fluorescence image (left image) of live cultured neurons expressing the α-syn-miniSOG-α-syn proteins. Following 4 minutes photooxidation, electron micrographs show labeled α-syn aggregates in cell bodies (middle image) as well as strong labeling in presynaptic terminals (right image).

**Supplementary Figure 15.**
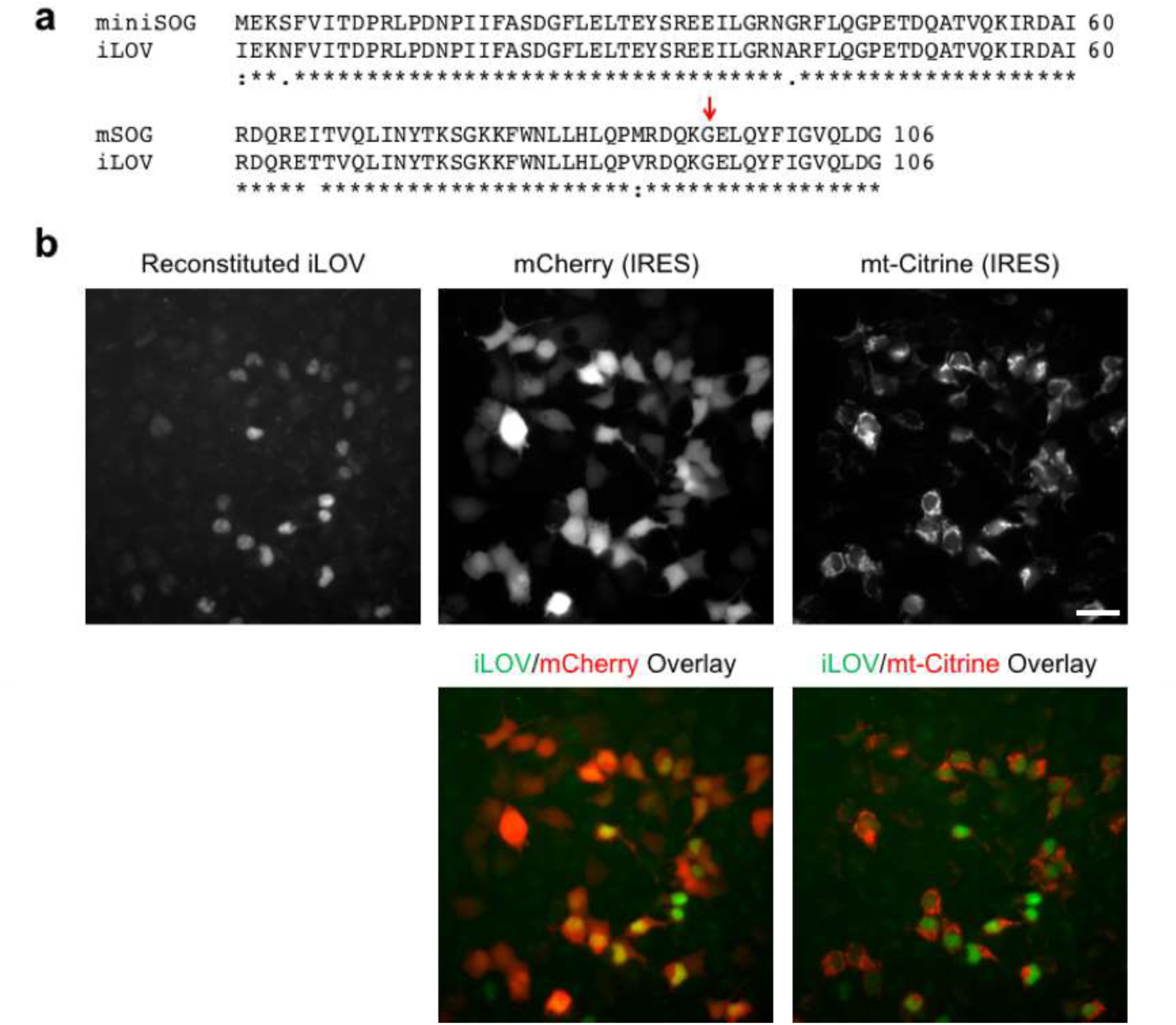
| The identified split-site is transferable to the related LOV domain and fluorescent reporter protein iLOV. (**a**) Sequence alignment of miniSOG and iLOV. The red arrow indicates the split position identified in this work. (**b**) Fluorescence images of HEK293 co-expressing bFos and bJun domains tagged with split iLOV fragments. Signal from reconstituted iLOV complexes were confined to nuclei, and fluorescence from IRES-mCherry and IRES-mtCitrine confirmed uptake of the two constructs. Scale bar, 30 microns.

**Supplementary Table 1.**
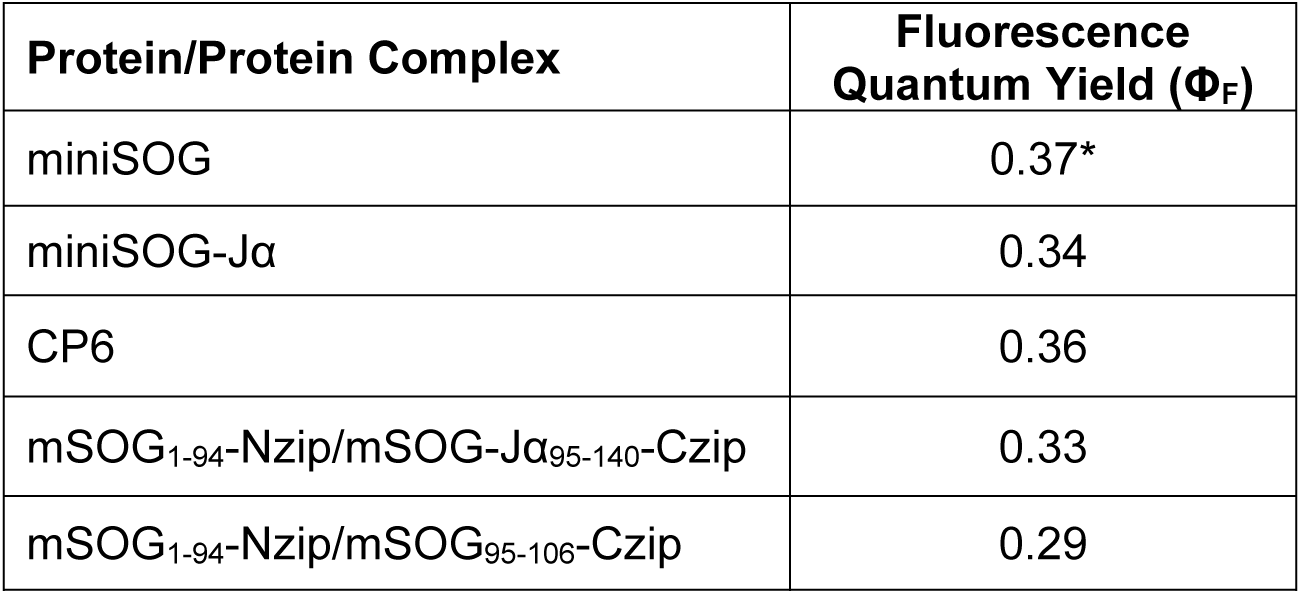
| Fluorescence quantum yields of the proteins and reconstituted complexes described in this study. Values were calculated using purified miniSOG as the standard (^∗^Shu, X., et al. *PLoS Biol*. **9**, e1001041 (2011)).

| Protein/Protein Complex | Fluorescence Quantum Yield ($\Phi_F$) |
| --- | --- |
| miniSOG | 0.37* |
| miniSOG-Jα | 0.34 |
| CP6 | 0.36 |
| mSOG<sub>1-94</sub>-Nzip/mSOG-Jα<sub>95-140</sub>-Czip | 0.33 |
| mSOG<sub>1-94</sub>-Nzip/mSOG<sub>95-106</sub>-Czip | 0.29 |

